# Inflammatory Signaling in Pancreatic Cancer Transfers Between a Single-cell RNA Sequencing Atlas and Co-Culture

**DOI:** 10.1101/2022.07.14.500096

**Authors:** Benedict Kinny-Köster, Samantha Guinn, Joseph A. Tandurella, Jacob T. Mitchell, Dimitrios N. Sidiropoulos, Melanie Loth, Melissa R. Lyman, Alexandra B. Pucsek, Toni T. Seppälä, Christopher Cherry, Reecha Suri, Haley Zlomke, Jin He, Christopher L. Wolfgang, Jun Yu, Lei Zheng, David P. Ryan, David T. Ting, Alec Kimmelman, Anuj Gupta, Ludmila Danilova, Jennifer H. Elisseeff, Laura D. Wood, Genevieve Stein-O’Brien, Luciane T. Kagohara, Elizabeth M. Jaffee, Richard A. Burkhart, Elana J. Fertig, Jacquelyn W. Zimmerman

**Affiliations:** Department of Surgery, Johns Hopkins University School of Medicine, Baltimore, MD; Department of Surgery, New York University Grossman School of Medicine, New York, NY; Department of Oncology, Sidney Kimmel Comprehensive Cancer Center, Johns Hopkins University School of Medicine, Baltimore, MD; Convergence Institute, Johns Hopkins University School of Medicine, Baltimore, MD; Bloomberg Kimmel Immunology Institute, Johns Hopkins University School of Medicine, Baltimore, MD; Department of Genetic Medicine, Johns Hopkins School of Medicine, Baltimore, MD; Faculty of Medicine and Health Technology, Tampere University and Tays Cancer Centre, Tampere University Hospital; Translational Tissue Engineering Center, Wilmer Eye Institute, Johns Hopkins School of Medicine, Baltimore, MD; Department of Biomedical Engineering, Johns Hopkins School of Medicine, Baltimore, MD; The Massachusetts General Hospital Cancer Center and Department of Medicine, Harvard Medical School, Boston, Massachusetts; Department of Radiation Oncology at New York University Grossman School of Medicine, NYU Langone Health, New York, New York; Department of Pathology, Johns Hopkins School of Medicine, Baltimore, MD; Department of Neuroscience, Johns Hopkins School of Medicine, Baltimore, MD; Department of Applied Mathematics and Statistics, Whiting School of Engineering, Johns Hopkins University, Baltimore, MD

**Keywords:** pancreatic ductal adenocarcinoma (PDAC), scRNAseq, organoids, cancer associated fibroblasts (CAFs), atlas

## Abstract

Pancreatic ductal adenocarcinoma (PDAC) is an aggressive malignancy characterized by a heterogeneous tumor microenvironment (TME) that is enriched with cancer associated fibroblasts (CAFs)^1^. Cell-cell interactions involving these CAFs promote an immunosuppressive phenotype with altered inflammatory gene expression. While single-cell transcriptomics provides a tool to dissect the complex intercellular pathways that regulate cancer-associated inflammation in human tumors, complementary experimental systems for mechanistic validation remain limited. This study integrated single-cell data from human tumors and novel organoid co-cultures to study the PDAC TME. We derived a comprehensive atlas of PDAC gene expression from six published human single-cell RNA sequencing (scRNA-seq) datasets^2–7^ to characterize intercellular signaling pathways between epithelial tumor cells and CAFs that regulate the inflammatory TME. Analysis of the epithelial cell compartment identified global gene expression pathways that modulate inflammatory signaling and are correlated with CAF composition. We then generated patient-derived organoid-CAF co-cultures to serve as a biological model of the cellular interactions learned from human tissue in the atlas. Transfer learning analysis to additional scRNA-seq data of this co-culture system and mechanistic experiments confirmed the epithelial response to fibroblast signaling. This bidirectional approach of complementary computational and *in vitro* applications provides a framework for future studies identifying important mechanisms of intercellular interactions in PDAC.

## Main Text

Pancreatic ductal adenocarcinoma (PDAC) remains challenging to effectively treat largely due to detection at advanced stages and a heterogeneous tumor microenvironment (TME) that limits treatment efficacy. Molecular changes in the epithelial cell population during carcinogenesis promote further changes in the surrounding non-epithelial cell populations and results in a dense and immunosuppressive TME^1^. The TME in PDAC is characterized by its heterogeneity and includes a variety of infiltrating cell types, including mesenchymal cells, such as cancer associated fibroblasts (CAFs), and multiple sub-populations of myeloid and lymphoid cells^1^. The recognition of phenotypic heterogeneity in cell types comprising each patient’s unique TME, and our understanding of the functional and dynamic diversity of these distinct cell populations, has historically been limited by available genomic technologies and represents a major barrier to our understanding of PDAC cancer biology.

Single-cell RNA sequencing (scRNA-seq) has recently enabled a more nuanced study of the PDAC TME, uncovering the heterogeneity in cell-type and function. Prior work with PDAC single-cell datasets has provided a roadmap to help identify individual cell populations and associated transcriptional regulation of the TME^8–11^. These data demonstrated previously underappreciated cellular heterogeneity throughout stages of malignant progression in humans and mouse models of PDAC tumors and are complemented by studies exploring the signaling pathways driving a tumor’s phenotype^12,13^. Single-cell technologies have furthered our capacity to identify discrete and functionally distinct subpopulations in both the malignant epithelial cells and stromal cells. CAFs in the PDAC TME are abundant and specific phenotypes have been implicated as either tumor enhancing or tumor-restraining subtypes^14,15^. Myofibroblastic CAFs (myCAFs) and inflammatory CAFs (iCAFs) are two subpopulations that have been well-described^4,16^. More recently, additional subtypes have been proposed, including those expressing major histocompatibility complex class II (MHC-II) genes, termed antigen-presenting CAFs (apCAFs)^4^. While the identification of these subpopulations and their associated characterization has opened new avenues of research in this disease, the mechanisms of intercellular interaction and specifically how CAF functional heterogeneity influences the phenotype of the tumor cells remains poorly understood.

To identify epithelial tumor cell and CAF intercellular interactions, we first collated an atlas of six published scRNA-seq datasets generated from small cohorts of PDAC patients^2–7^. Single-cell data provide opportunities to isolate subpopulations for inferring intercellular interactions and identifying patterns of cell signaling associated with individual cell types or cell stressors. To investigate the inflammatory signaling responsible for major histocompatibility class (MHC) II expression in tumor cells demonstrated in prior work^17–19^, we focused on inflammatory signaling in epithelial tumor cells learned from applying our Bayesian non-negative matrix factorization method CoGAPS^20^ for discovery of gene expression patterns in this comprehensive atlas. Even with a larger cohort for analysis, *in silico* analyses of these data are limited in their ability to evaluate mechanism and implications of intercellular interactions. Thus, innovations in laboratory systems and complementary computational approaches are needed to accelerate further the study of intercellular interactions and enable *in vitro* mechanistic evaluation of *in silico* hypotheses generated from high-throughput datasets.

Currently, available biological models of PDAC are restricted in large part to either mouse models or *in vitro* epithelial cell lines derived from human pancreatic tumor cells. While mouse models can provide a global characterization of the TME, depletion experiments are required to isolate the effect of individual cell types, and this approach is limited by a lack of depleting agents with specificity for functionally different TME cell subtypes. *In vitro* experiments have traditionally been limited to epithelial tumor cells, but more recently have been adapted to investigate the effect of cell-cell interactions and paracrine crosstalk with other cell types using conditioned media. However, organoids and organoid co-cultures are an emerging system that may best recapitulate the tumor and associated microenvironment^21^.

To further our ability to dynamically examine intercellular interactions inferred from single-cell analysis, we established an *in vitro* three-dimensional co-culture system of patient-derived organoids (PDOs) and patient-derived CAFs. This system is useful for evaluating the transcriptional dynamics in response to external stimuli and investigating core intercellular interactions. Combining this approach with our novel suite of computational tools for single-cell analysis enables bi-directional investigation of fibroblast-tumor cell interactions between human tissue and PDO biological models, which is broadly applicable to untangling the complexities of intercellular interactions in the PDAC TME.

### A harmonized atlas of gene expression in PDAC created from an integrative analysis of 6 single-cell data to explore inflammatory signaling pathways

To explore inflammatory signaling between CAFs and tumor cells in PDAC, we integrated six published human scRNA-seq datasets into a single comprehensive atlas. In total, the atlas reflected gene expression in 174,394 total cells from 61 PDAC (142,807 cells) and 16 non-malignant pancreatic tissue samples (31,587 cells) **(Figure 1A-B, Supplemental Figure 1** with subset of PDAC specimens)^2–7^. All samples were of pancreatic origin (no metastatic samples) and from treatment-naïve patients **(Table 1)**. Of the 61 PDAC samples (2 were described as arising from cystic pancreatic lesions), 52 samples originated from patients with apparent localized disease, 6 samples originated from patients with distant metastases, and 3 samples were from patients whose stage was unknown. Of the 16 non-malignant control samples, 5 were specified by the authors as normal-adjacent to adenocarcinoma and 11 were derived from samples described as normal-adjacent to non-malignant pathologies. All control samples and the majority of PDAC samples were obtained from resected surgical specimens. Ten PDAC samples originated from fine-needle biopsies, four of which presented with clinical metastatic disease.

**Table 1.**
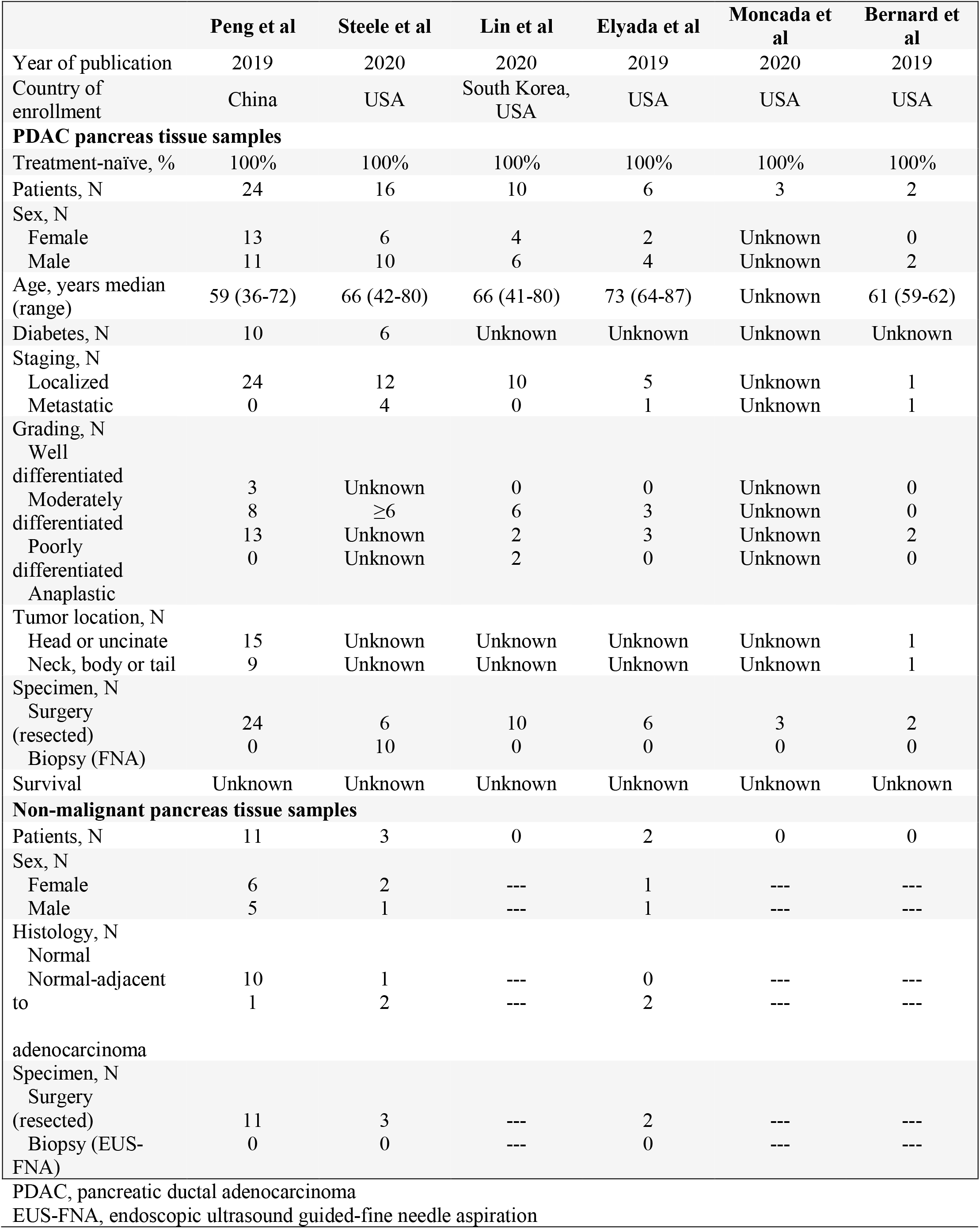
Clinical patient data extracted from the six harmonized published datasets integrated in the PDAC atlas.

**Figure 1.**
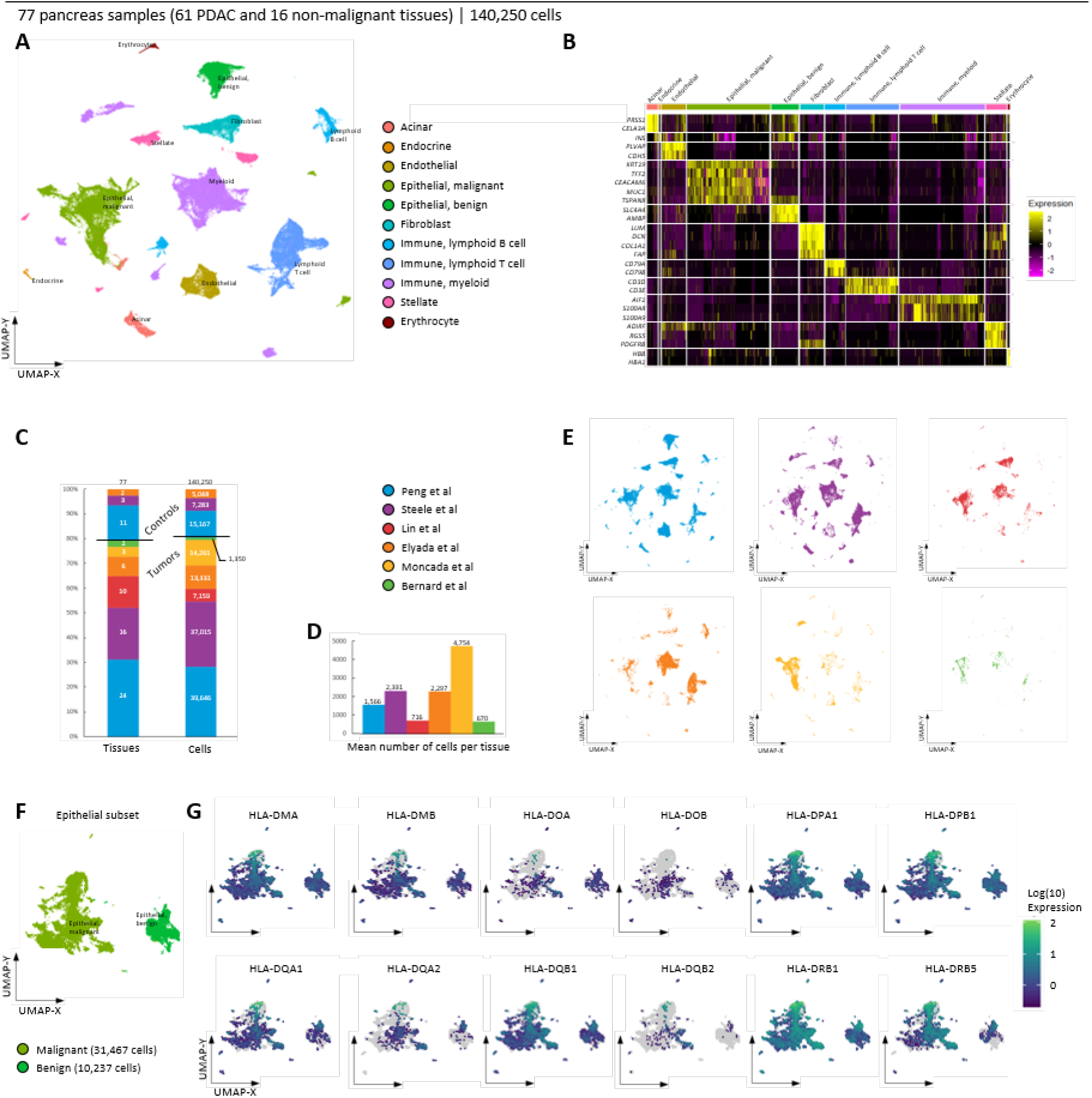
Summary of atlas composition and evaluation of MHC-II gene expression in atlas samples. (A) Complete atlas with assigned cell types. (B) Heatmap of differentially expressed genes used for cell type annotations. (C) Relative contribution of the 77 different samples with 140,250 cells, separated by tumor (below line) and control tissue (above line). (D) Mean number of cells per tissue by dataset origin. (E) Cell mapping by dataset origin from the six manuscripts in the complete atlas. (F) Epithelial cell clusters from patient tumor samples. (G) Expression of 12 MHC-II genes was queried in the atlas demonstrating variable endogenous expression of each gene. UMAP: Uniform Manifold Approximation and Projection.

After filtering cells based upon biological or technical quality metrics (fraction of mitochondrial reads and number of transcribed genes as defined in the methods), the resulting atlas included 140,250 cells (dropout 19.6%). The median and mean cell counts per patient sample were 1,455 and 1,821, respectively (interquartile range 828 – 2,200). Following computational pre-processing, we performed a clustering analysis for cell type annotation (supplemental methods). To make inferences on biology from an integrated atlas, it is critical to first determine inter-dataset variation for the mitigation of non-representative findings and technical artifacts. Most of the contributing tissues and cells (35 and 54,813, respectively) originated from Peng et al^3^ **(Figure 1C-E)** with a mean of 1,566 cells per tissue. The mean number of included cells per sample was highest (4,754 cells) in the dataset by Moncada et al^5^, a set that contributed PDAC samples from only three patients **(Figure 1C-E)**. Despite the heterogeneity in cell count, mapping dataset of origin on the atlas demonstrated contribution from all six datasets for most clusters, thereby visually verifying our integration strategy **(Figure 1E)**.

Within PDAC samples, the predominant cell populations were epithelial malignant cells (32,515 or 29.3%), cells of myeloid origin (28,971 cells or 26.1%), and T cells (17,284 cells or 15.6%) **(Supplemental Figure 1B)**. The mesenchymal cell populations were composed of 8,953 CAFs (8.1%) and 6,049 stellate cells (5.4%). Altogether, the components of the TME (immune and mesenchymal cell populations combined) contributed 67,690 cells or 60% of the total cell count. In the myeloid and lymphoid cell clusters, we derived subpopulations reflecting macrophages, mast cells, neutrophils, regulatory T cells and natural killer/cytotoxic T cells **(Supplemental Figure 2)**. For the epithelial cells from tumor tissues, copy number variation analyses (CNV, **Supplemental Figure 3**) were combined with differential gene expression analyses to confirm epithelial cell classification (benign or malignant). In the epithelial cell cluster, we classified malignant cells from PDAC tissues according to the classical and basal gene expression programs first reported by Moffitt et al^21^ **(Supplemental Figure 4)**. In the CAF cluster, cells were subtyped according to iCAF and myCAF gene expression programs **(Supplemental Figure 5)**. Cell cycle analyses within the PDAC samples revealed high cell cycle activity reflected by increased activity scores and translated phases primarily in the epithelial malignant, acinar, myeloid and CAF populations **(Supplemental Figure 1C-D)**.

To evaluate MHC-II expression in the PDAC epithelial compartment, we queried the atlas for expression of 12 MHC-II genes, including HLA-DRB1, HLA-DOA, HLA-DMA, and identified heterogenous expression in the tumor epithelial cell population **(Figure 1F-G)**. Overall, MHC-II gene expression was most pronounced in epithelial cells derived from tumor tissue as compared to epithelial cells obtained from non-malignant, control samples **(Supplemental Figure 6)**. Additionally, strong expression of HLA-DO and HLA-DM isoforms suggest functionality of MHC-II proteins given their role as chaperones in antigen presentation.

### Inflammatory signaling drives MHC-II expression in the epithelial compartment of PDAC

Next, we investigated patterns of gene expression within the epithelial cell populations in the atlas. To distinguish epithelial cells within PDAC tissues from epithelial cells derived from non-malignant samples, we performed an unsupervised analysis of gene expression patterns with our single-cell Bayesian non-negative matrix factorization algorithm CoGAPS^22^. Batch effects were mitigated by identifying robust gene expression patterns that are maintained across the two largest sample cohorts that contain both PDAC and non-malignant samples: Peng et al^3^ (18,261 epithelial cells) and Steele et al^2^ (7,181 epithelial cells). These two datasets account for 61.0% of all epithelial cells in the atlas. Mapping the epithelial cells from these two datasets on the UMAP from the entire atlas confirmed that they represent 49.3% of all malignant cells, and 97.1% of all benign cells. We then applied the CoGAPS algorithm to interrogate patterns of gene expression across these epithelial populations, identifying 8 distinct patterns **(Supplemental Figure 7)**. Each pattern was annotated by estimated overrepresentation across genes identified by the CoGAPS pattern marker statistic^23^ and Hallmark gene sets from the Molecular Signatures Database^24^. Pattern 1 identified pathways associated with UV response and TGFβ signaling that did not reach statistical significance **(Supplemental Table 1)**. Pattern 2 identified pathways of estrogen response and KRAS signaling that are relevant for PDAC development. Metabolic pathways including cell cycle, oxidative phosphorylation, and glycolysis were significant in Patterns 3-5. Pattern 6 identified pathways of apoptosis activity and Pattern 8 was dominated by genes inherent in response to hypoxia. Notably, genes associated with Pattern 7 were classified by CoGAPS to included increased pathway activities in inflammatory, fibrogenic, and malignant progression-associated gene sets, including epithelial-mesenchymal transition (EMT) **(Figure 2A-C)**.

**Figure 2.**
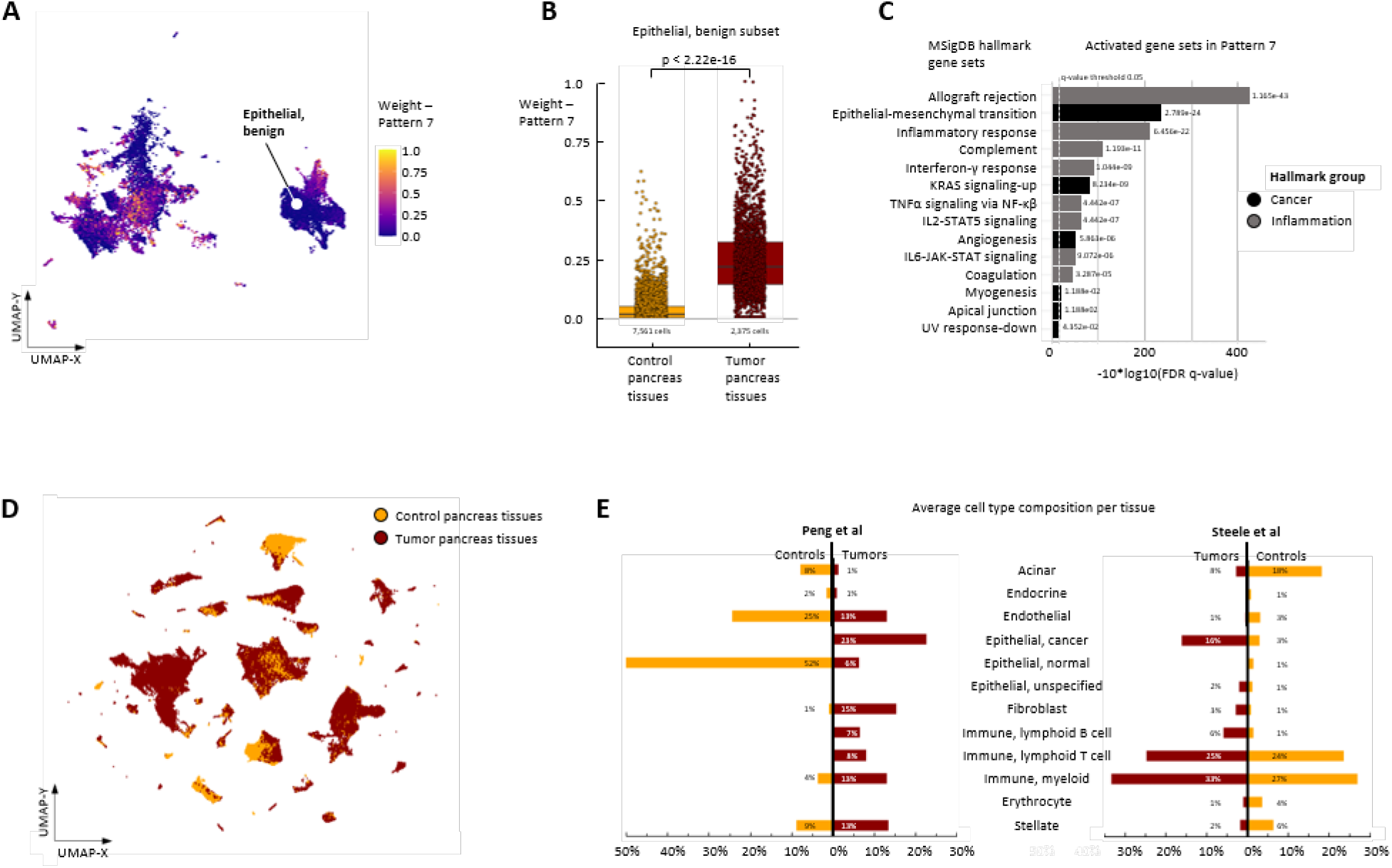
Identification of pattern of inflammation (Pattern 7) in the atlas and cell type distribution in the atlas. (A) Complete atlas subset of the epithelial cell populations (malignant, benign and unspecified) from the Peng et al^3^ and Steele et al^2^ data with assigned weights of Pattern 7 as resulting from CoGAPS analyses. (B) Boxplot of Pattern 7 weights within the Epithelial, benign cell population demonstrating differences between control and tumor pancreas tissues. p < 2.22 e-16, generated by Wilcoxon test. (C) Overrepresented MSigDB hallmark gene sets in cells expressing Pattern 7 genes. (D) Cell mapping by tumor vs. control pancreas tissue in the complete atlas. (E) Differences in cell type composition for selected tumor vs. control pancreas tissues originating from Peng et al^3^ and Steele et al^2^.

Annotation of the genes associated with Pattern 7 using the hallmark gene sets as a guide demonstrated enrichment in this pattern for processes of inflammation in the epithelial compartment **(Figure 2C)**. We also observed that Pattern 7 weights were significantly higher in both the malignant epithelial cells and the benign epithelial cells derived from patients with tumors relative to the epithelial cells in the non-malignant control samples **(Figure 2B)**. The enrichment of Pattern 7 within the malignant epithelial population suggests a potentially transformative inflammatory program drives phenotype in the epithelial compartment in a manner similar to that which has been described in non-epithelial cell types in PDAC^25^. These findings are further supported by inducible MHC-II expression in response to inflammatory stimuli anticipated in the malignant epithelial compartment. We note that the MHC-II expression is identified in the malignant epithelial cell compartment in a manner independent of the specific inflammatory stimuli defined by Pattern 7, suggesting additional mechanisms driving MHC-II activation in epithelial cells may be playing a role.

To investigate alternative sources of activated inflammatory pathways in the TME, we next investigated the fibroblast compartment in the atlas. Fibroblasts are an important driver of inflammatory processes in many human diseases, though much of this association has been established in non-cancer inflammatory illnesses^26^. Therefore, we hypothesized that CAFs may also drive inflammatory signaling in the TME that influences malignant epithelial cells. To test this hypothesis, we annotated the cell type composition in the tumor and control samples to identify subpopulations in the TME. Notably, the non-epithelial populations in the atlas largely originated from tumor samples **(Figure 2D)**. When comparing the average cell type composition between the control and tumor samples within the Peng et al^3^ dataset, the fraction of fibroblast, myeloid and lymphoid populations were greater in tumor samples, with a proportional decrease in cells of endothelial origin, acinar cells and total epithelial cells **(Figure 2E)**. Within the Steele et al^2^ dataset, differences between the fibroblast/CAF, myeloid and lymphoid populations, while present, were less pronounced **(Figure 2E)**. To elucidate the impact of CAF signaling in the TME on epithelial inflammatory gene expression, we correlated the mean CoGAPS weights of the inflammatory pattern 7 in the epithelial compartment with the presence of CAFs in the TME using the datasets from Peng et al^3^ and Steele et al^2^. We identified a direct association between increasing fibroblast proportion in the TME and the mean weight of CoGAPS Pattern 7 in the epithelial cells **(Supplemental Figure 8)**. This association was lost when fibroblast populations were further divided into iCAFs and myCAFs **(Supplemental Figure 8)**. Therefore, uncovering the mechanisms that drive CAF-mediated inflammatory signaling in the epithelial compartment requires further validation using approaches beyond the publicly-available gene expression datasets.

### PDOs as a system to evaluate mechanisms of inflammatory signaling in PDAC: Epithelial cells respond to interferon gamma (IFNγ) stimulation over time to induce MHC-II

We further utilized our PDO cultures to validate the inflammatory phenotype identified in the atlas and to interrogate the mechanisms of inflammatory gene expression in the setting of multi-compartmental three-dimensional co-cultures. We previously demonstrated that PDAC PDOs, comprised exclusively of malignant epithelial cells, have the capacity to recapitulate intratumoral heterogeneity and accurately provide clinically-actionable data for chemotherapeutic selection^27,28^. Previous work has also shown that 24 hours of 200ng/mL of IFNγ added into cell media induces MHC-I and PD-L1 upregulation in organoids^29^. To extend these findings in additional samples from our organoid bank, we screened eleven PDO lines by flow cytometry (gating strategy **Supplemental Figure 9**) for both constitutive and IFNγ-induced cell surface protein expression of MHC-I and II and PD-L1 **(Figure 3A)**. At 24 hours, both MHC-I and PD-L1 demonstrated a robust increase in expression in response to IFNγ stimulation. HLA-DR was used as a representative marker for MHC-II expression in this system. As expected, HLA-DR expression was restricted at baseline and remained low during the first 24 hours of IFNγ stimulation. Longer-term exposure to IFNγ treatment demonstrated upregulation of HLA-DR expression over 96 hours consistent with induced gene expression changes to MHC-II as a result of inflammatory signaling **(Figure 3B)**. Reproducibility of this phenotype was demonstrated in a set of PDOs with evaluation and characterization of MHC-II presence in the setting of both endogenous and IFNγ-induced gene expression changes **(Figure 3C-D)**. To further evaluate IFNγ-induced changes in MHC-II alleles which lack specific antibodies, we validated HLA-DRB1, HLA-DRA, HLA-DPB1, HLA-DQB1 by qPCR and observed a heterogenous IFNγ-induced response in gene expression **(Figure 3E)**. To further confirm these data, we examined inducible MHC-II expression in formalin fixed and paraffin-embedded (FFPE) PDAC PDOs using IHC with antibodies against HLA-DR and HLA-DR/DP/DQ to inform the spatial distribution of the MHC-II expression across the PDO system **(Figure 3F-G)**, further confirming robust MHC-II upregulation in response to IFNγ.

**Figure 3.**
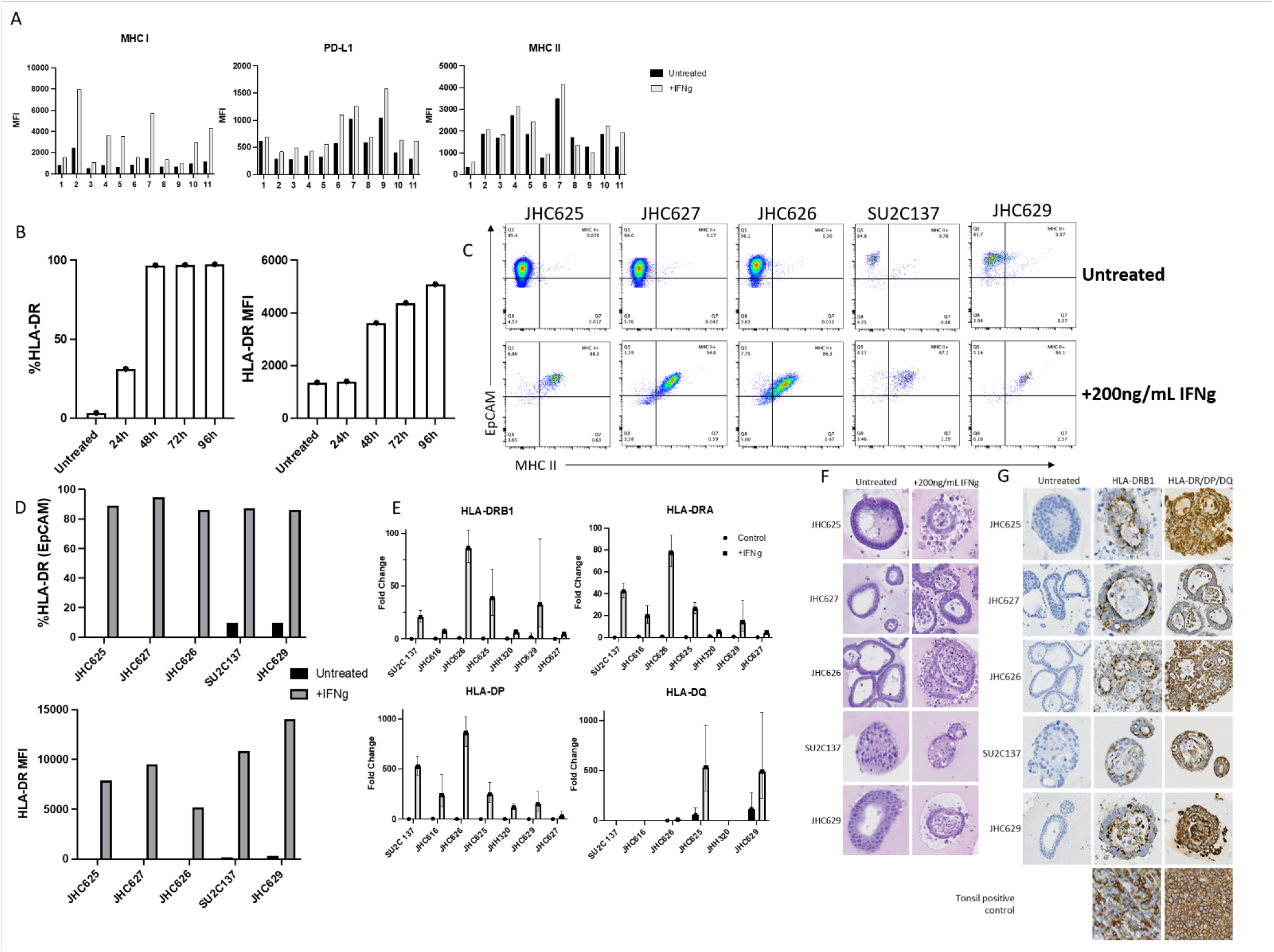
MHC-II expression in PDAC PDOs. (A) Cell surface MHC-I, PD-L1, and MHC-II expression as determined by flow cytometry. Organoids were stimulated with 200ng/mL of IFNγ for 24hr or were left unstimulated. Bar graphs represent Mean Fluorescence Intensity (MFI). (B) HLA-DR (MHC-II) expression as determined by flow cytometry is increased after IFNγ stimulation for 24-96 hours shown both by % increase (left panel) and by increase in MFI (right panel) across one PDO line. (C) Representative flow cytometry dot plots of 5 organoid lines that were treated with 200ng/mL of IFNγ for 96 hours or left unstimulated. (D) Bar graph quantification of plots in C, bar graphs represent % increase in HLA-DR (MHC-II) expression of EpCAM+ cells (top panel) and Mean Fluorescence Intensity (MFI) (bottom panel). (E) qPCR readout of organoid lines that were stimulated with 200ng/mL of IFNγ for 96 hours or left unstimulated; genes quantified include HLA-DRA/HLA-DRB1/HLA-DPB1/HLA-DQB1. P-value for HLA-DRA/HLA-DRB/HLA-DP when tested between control and treated groups = 0.0006, p-value for HLADQ = 0.4 (statistical analysis was performed using a two-tailed, unpaired, parametric Mann-Whitney test). (F) H&E slides of PDOs with and without IFNγ induction. Organoids were harvested 96 hours after IFNγ induction. (G) Immunohistochemistry (IHC) demonstrating expression of HLA-DRB1 (middle) and antibody binding to the common beta chain of HLA-DR/DP/DQ (right) after 96 hours of IFNγ stimulation.

### Co-culturing PDOs and CAFs enhances the epithelial inflammatory pattern identified in atlas samples and drives expression of MHC-II genes

To evaluate the mechanisms responsible for fibroblast-induced inflammatory signaling towards the malignant epithelial compartment that was identified in the atlas, we established a system of PDAC PDOs co-cultured with patient-derived CAFs. Each patient-matched PDO-CAF co-culture leverages our previously published methods for generating PDOs^27,28^ while concurrently extracting CAFs from surgical resection specimens **(Supplemental Figure 10)**. The three-dimensional cultures were established in Matrigel as the surrogate basement membrane for suspending culture components. We then optimized the culture conditions to maximize both the viability of input cell types during culture and the viability of cells following extraction **(Supplemental Figure 10)**. These methods for co-culture maintain viability of both cell-types allowing for mechanistic study **(Figure 4A)**.

**Figure 4.**
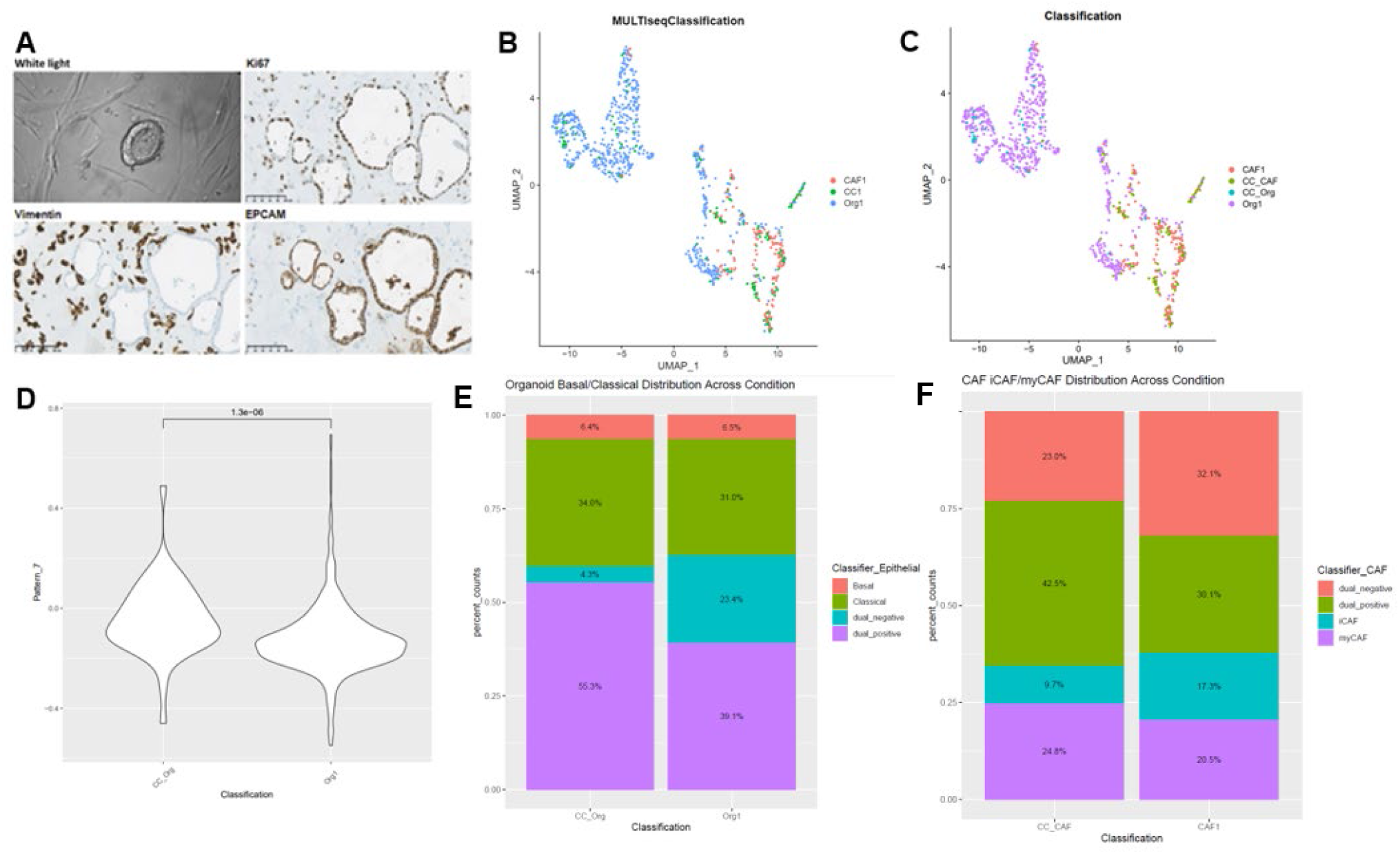
Patient-derived organoids co-cultured with CAFs recapitulate the inflammatory pattern identified in tumor epithelial cells and demonstrate dynamic cellular phenotypes. (A) Representative brightfield image of co-culture at 20x magnification. Representative IHC of co-culture demonstrating proliferation by Ki-67 after co-culture (top right), vimentin positive CAFs (bottom left), and EpCAM positive organoids (bottom right). Images obtained at 20x magnification. (B) UMAP demonstrating culture conditions: organoid monoculture (Org1), CAF monoculture (CAF1), co-culture (CC1). (C) UMAP demonstrating cell-type calls after co-culture: Organoid monoculture (Org1), CAF monoculture (CAF1), CAFs from co-culture (CC_CAF), organoids from co-culture (CC_Org). (D) Pattern 7 is composed of inflammatory genes and was enhanced in organoid cells from co-culture relative to organoid cells from monoculture, p=1.3e-6. (E) Co-culture demonstrates dynamic epithelial representation in the co-culture condition with a greater percentage of cells representing both basal and classical markers (dual_positive) present in co-culture. (F) Co-culture demonstrates dynamic CAF representation in the co-culture condition with a greater percentage of cells representing both iCAF and myCAF markers (dual_positive) present in co-culture.

To compare the signaling processes in our co-culture to the inflammatory signals observed in human PDAC from our atlas, we performed scRNA-seq profiling during a 12-hour PDO-CAF co-culture. Controls included scRNA-seq for both PDO and CAF monoculture. Multiplex analysis, with the established MULTI-seq protocol^30^, was used to assess transcriptional heterogeneity in the epithelial and CAF compartments. As expected, clustering analysis and visualization of the scRNA-seq data from these conditions separated the fibroblasts from epithelial cells. Our initial evaluation demonstrated that cells from the co-culture and monoculture conditions did not separate into distinct clusters **(Figure 4B-C)**. Despite the lack of separate clusters from the co-culture-derived cells, we did identify transcriptional plasticity in the epithelial compartment with the co-culture condition demonstrating a greater number of cells expressing markers for both the classical and basal epithelial subtypes and fewer cells lacking expression consistent with either classification **(Figure. 4E)**^31^. Baseline heterogeneity of the basal and classical programs in PDOs identified using deconvolution of bulk RNA-seq was consistent with the recent report by Krieger et al^32^ **(Supplemental Figure 11)**. Moreover, induced plasticity was not limited to the epithelial compartment as the co-culture also demonstrated increasing proportions of CAFs expressing gene markers for both iCAFs and myCAFs **(Figure 4F)**. This further supports the notion that co-culture induces both epithelial and CAF transcriptional plasticity.

We hypothesized that the CAFs in this organoid co-culture drive inflammatory signaling in a manner similar to that observed in the human system. Therefore, we sought an integrative analysis to quantify the similarity between the scRNA-seq datasets from our human atlas and PDO co-culture. This analysis was performed using our transfer learning method ProjectR^33^ to project the inflammatory pattern (CoGAPS Pattern 7), identified in the epithelial cells from the human PDAC atlas, onto the epithelial cells of the PDO-CAF co-culture. After 12 hours of co-culture, we identified increased gene expression in the Pattern 7 gene signature in PDAC organoids co-cultured with CAFs, relative to those cultured in mono-culture **(Figure 4D)**. These data further support the hypothesis that the inflammatory signaling observed in epithelial cells is stimulated by the presence of CAFs in the TME.

We next examined CAF-mediated upregulation of MHC-II in malignant epithelial cells by characterizing MHC-II expression from the co-culture scRNA-seq data **(Figure. 5A-C)**. Similar to data seen under IFNγ stimulation, baseline and early (12 hours) HLA-DRA and HLA-DRB expression was low in both the monoculture and co-culture conditions **(Figure 5 A-C)**. “To expand beyond 12 hours and better understand if there is a temporal change in the epithelial compartment over a longer period, we co-cultured PDO-CAF for 24 and 96 hours. At each timepoint, we flow sorted the co-culture to have a pure population of cells to query MHC-II gene expression changes by qPCR. Similar as our findings with IFNg treatment, HLA-DRA and HLA-DRB expression increased with increasing time in co-culture conditions up to 96 hours **(Figure 5D-E)**. This further implicates CAFs as critical mediators of epithelial inflammatory signaling in the tumor microenvironment.

**Figure 5.**
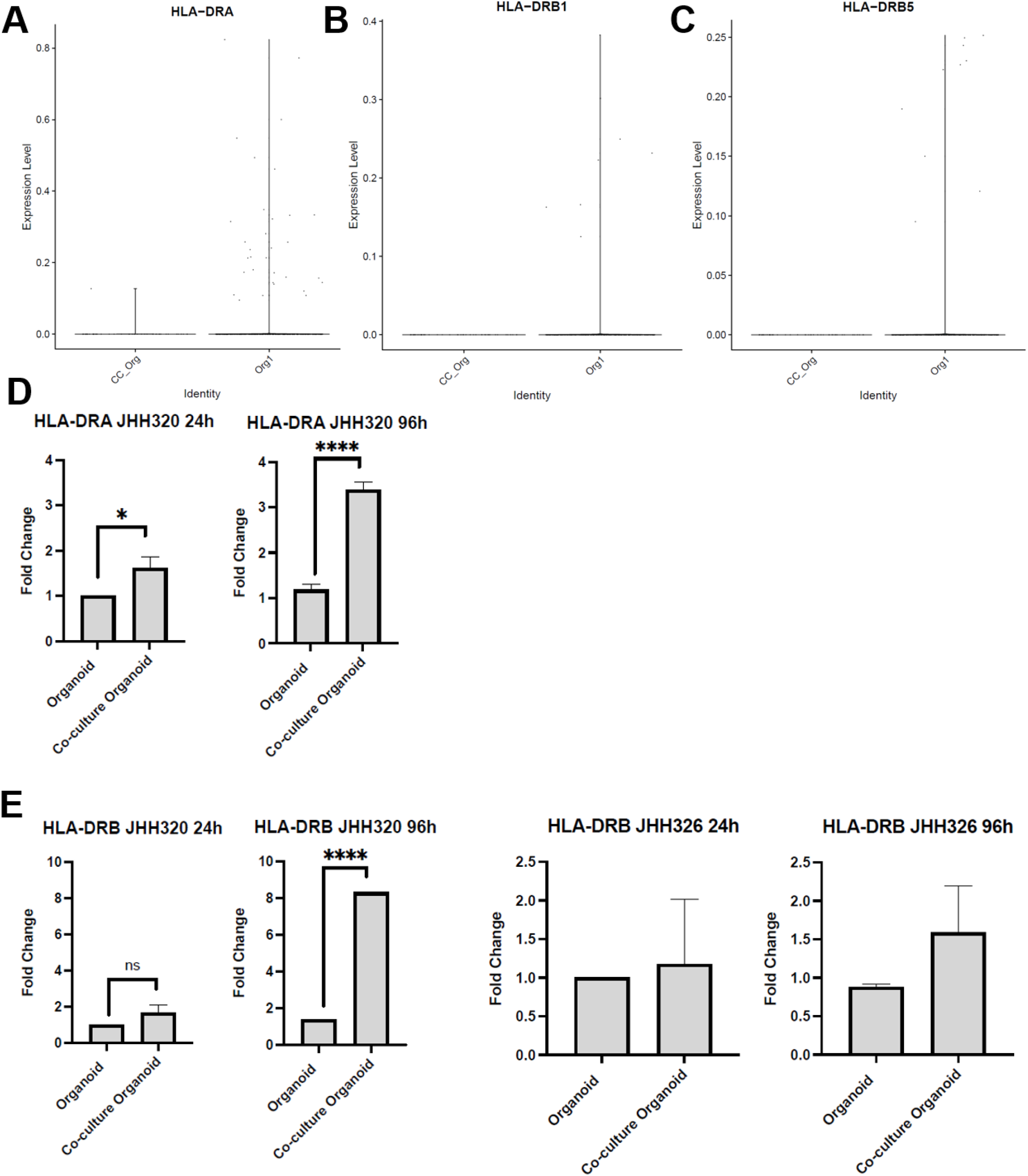
Evaluation of MHC-II expression in the co-culture in response to the presence of CAFs. Expression of (A) HLA-DRA, (B) HLA-DRB1, and (C) HLA-DRB5 after co-culture for 12 hours evaluated using MULTI-seq. Expression is limited in all populations. (D) HLA-DRA and (E) HLA-DRB expression after 24 and 96 hours of co-culture after which cells were flow sorted prior to qPCR. HLA-DRA did not amplify in JHH 326 at either 24 or 96 hours. HLA-DRB demonstrated less consistent amplification limiting statistical analysis. Co-cultures were established from two patients for which there were matched PDOs and CAFs. Plotted are the Fold Change values comparing our PDO co-culture to monoculture using GAPDH as an endogenous control. Comparisons of monoculture and co-culture conditions are statistically supported using the two-tailed students t-test with equal variance in PRISM (V9.2.0 [283]). Significance is measured as: ****, p<0.0001; ***, p<0.001; **, p<0.01; *, p<0.05; ns, not significant.

### Domino analysis of intercellular signaling identifies VEGF-A as an epithelial-derived ligand and ITGB1 as a fibroblast-derived ligand driving molecular signaling between fibroblasts and malignant epithelial cells

Next, we performed bidirectional *in silico* and *in vitro* experimental interrogation of the mechanisms leading to crosstalk between CAFs and epithelial cells that underlies the observed Pattern 7 inflammatory signaling in PDAC. No consistent association was observed between the composition of epithelial and CAF subtypes across the cohort of PDAC tumors in the atlas **(Supplemental Figure 12)**. Still, we hypothesized that signaling pathways underlie epithelial and CAF interactions that are consistent between human tumors and our organoid co-culture model. To infer these pathways, we selected the epithelial and CAF populations of the PDAC tumor atlas derived from the Peng et al^3^ (12,120 epithelial cells, 63.8%; 5,823 CAFs, 84.0%) and Steele et al^2^ (6,883 epithelial cells, 36.2%; 1,110 CAFs, 16%) datasets. We then analyzed these data with Domino, a computational method that infers intercellular interactions in scRNA-seq data by quantifying for coordinated gene expression changes between the ligands of one cell type with receptors of another^34^. The resulting analysis generated a putative global signaling network in PDAC based on population-specific gene expression of ligands and receptors with established signaling relationships **(Figure 6A)**. Across both datasets, *VEGF-A* was expressed by epithelial populations and predicted to target fibroblasts **(Figure 6B-C)**. To validate this inference in human tissue, we set about examining this relationship in our PDO-CAF co-culture. Cells were flow sorted after 24 or 96 hours of co-culture and qPCR was performed to assess *VEGF-A* expression in the co-culture as compared to the monoculture. At both timepoints, *VEGF-A* demonstrated increased expression in cells derived from the co-culture relative to those extracted from monoculture, consistent with the Domino inference **(Figure 6D)**.

**Figure 6.**
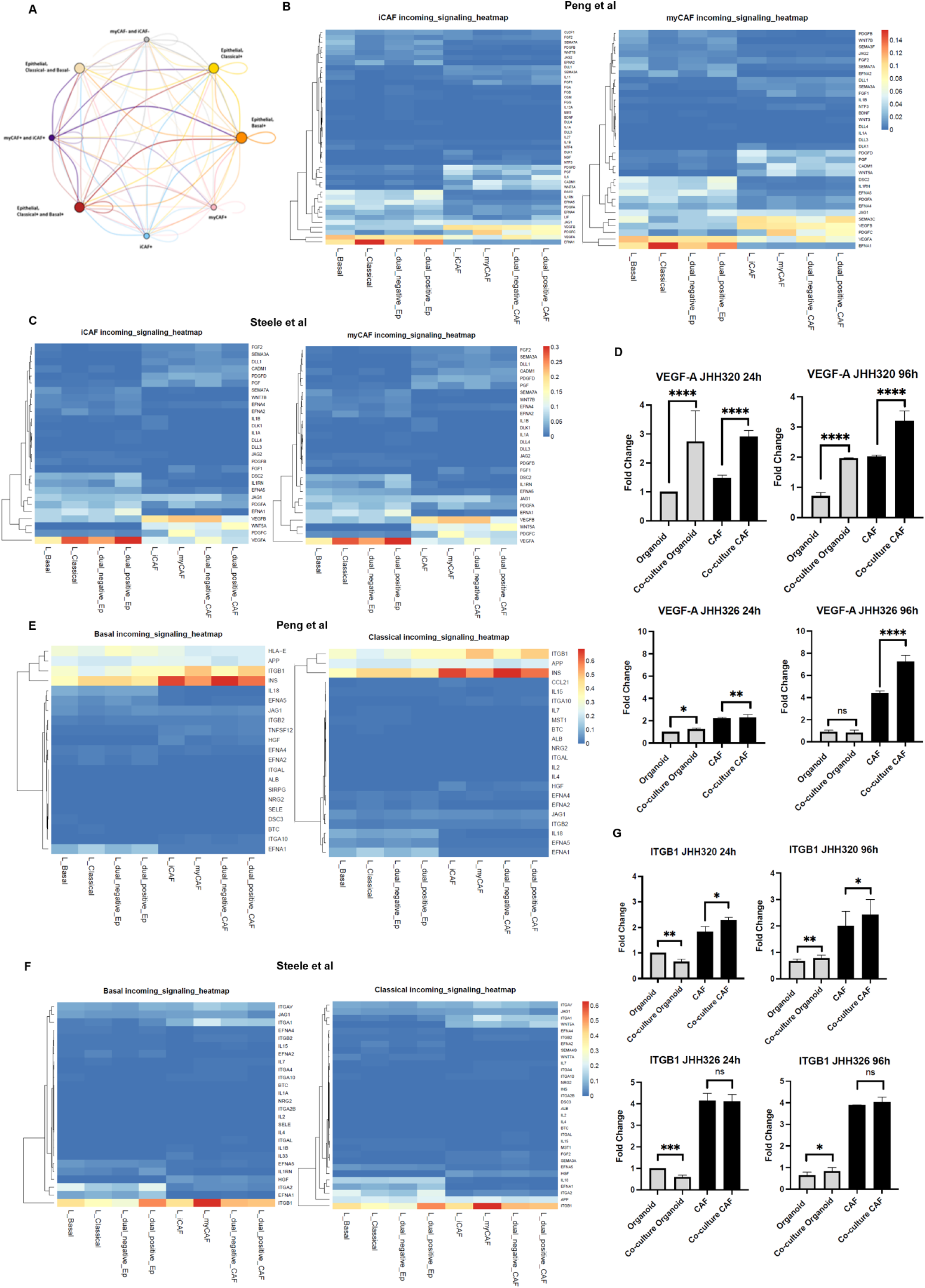
Domino evaluation of intercellular interactions in the atlas with validation in PDO-CAF co-culture. (A) Signaling network between epithelial and CAF subpopulations from tumor pancreas tissues in the Peng et al^3^ dataset as derived from the Domino R package. Nodes of the subpopulations are sized according to the amounts of expressed targeting ligands. The thicknesses of the intercellular connections are scaled based on the strength of signaling with their color indicating the signals’ origin (directionality). (B) Heatmap demonstrating *VEGF-A* as a ligand originating in the epithelial populations with the CAFs receiving this signal from the Peng^3^ and (C) Steele^2^ datasets. (D) Differential expression by qPCR of *VEGF-A* in monoculture and co-culture CAFs and epithelial cells after 24 or 96 hours of co-culture. (E) Heatmap demonstrating *ITGB1* as a ligand originating in the CAF populations with the epithelial cells receiving this signal from the Peng et al^3^ and (F) Steele et al^2^ datasets. (G) *ITGB1* expression in monoculture and co-culture CAFs and epithelial cells after 24 or 96 hours of co-culture. Plotted are the Fold Change values comparing our PDO co-culture to monoculture using GAPDH as an endogenous control. Comparisons of monoculture and co-culture conditions are statistically supported using the two-tailed students t-test with equal variance in PRISM (V9.2.0 [283]). Significance is measured as: ****, p<0.0001; ***, p<0.001; **, p<0.01; *, p<0.05; ns, not significant.

Finally, we evaluated intercellular signaling, derived from CAFs, that drives gene expression in epithelial PDOs. Across basal and classical epithelial subtypes, *ITGB1* was identified in both the Peng et al^3^ and Steele et al^2^ datasets as a ligand from both iCAFs and myCAFs directed at the epithelial subpopulations **(Figure 6E-F)**. While *ITGB1* expression was identified both in patient-derived organoids and CAFs in our co-culture model, it was differentially expressed in the fibroblasts **(Figure 6G)**. Although *ITGB1* expression is not tissue- or cell-type specific, its expression is classically associated with fibroblasts and impact the structure and function of the extracellular matrix. Taken together, discovery of this novel mechanism of CAF-epithelial cell inflammatory signaling, which was inferred by computational approaches using human tumor data and then validated in PDO-CAF co-culture, we demonstrate the utility of integrating human scRNA-seq data from human tissues with co-culture models to provide bidirectional study of critical TME effects on the cancer cells in PDAC tumors.

## Discussion

Despite improving outcomes and new therapeutic regimens for the majority cancer subtypes, PDAC outcomes have remained stubbornly resistant to novel targeted therapeutics and new treatment approaches. Recent technologic advances have shown that PDACs are composed of a complex and heterogenous TME that actively restricts therapeutic access and limits cancer cell killing. In addition, studies suggest that cancer cells co-opt normal stromal cell behavior, inducing fibroblasts to become CAFs that support PDAC development and progression^35^. Although the genomic landscape has identified oncogenes associated with this process in PDAC, our understanding of additional mechanisms contributing to tumorigenesis is limited by a lack of characterization of the complex TME. Single-cell technologies, and the computational tools used for their analysis, enhance our ability to characterize the impact that this unique TME can impart on structure and function of malignant epithelial cells in PDAC. These technologies have empowered unprecedented discovery of cellular function directly in human studies. Nevertheless, one persistent barrier to advancement in the field is that there are limited relevant experimental systems with a capacity to mimic the human tumor TME and thereby enable mechanistic study to determine the function of *in silico* predictions in single-cell analysis. Here we present a convergence approach to overcome this barrier, using a novel PDO-CAF coculture system to validate changes in signaling between cells in the TME that we have combined with *in silico* inference through transfer learning from single-cell analysis of publicly available datasets.

This study combined an *in silico* analysis of transcriptional states in human tissue using our novel suite of computational tools with PDO-CAF co-culture to examine the role of inflammatory gene expression patterns in PDAC. This approach was facilitated by the evaluation of dynamic intercellular interactions using a single-cell atlas to uncover cell-specific expression patterns that were then validated in cell-specific organoid co-cultures from additional patient tumors. Notably, the inflammatory gene expression patterns observed in the single-cell data from human tumors were also observed in tumor-adjacent epithelial cells that were transcriptionally more suggestive of “benign” epithelium. These findings provide additional data to support earlier observations that the tissue surrounding a tumor, even that in the epithelial compartment, is phenotypically distinct from normal epithelial cells obtained from a non-malignant specimen^36^. Consistent with this observation, our complementary study of spatial transcriptomic profiling in human pancreatic intraepithelial neoplasia (PanIN) demonstrates dynamic inflammatory signaling in epithelial cells during tumor progression^37^. These data provide a model in which the changes to epithelial cells induced by the biological alterations in the background TME contribute to subsequent tumorigenicity of the malignant epithelial state, with a transition between inflammatory and growth signaling that is further supported in our complementary PanIN data.

Previous studies have shown that tumor cells from numerous cancer types can be induced to express MHC-II, but less is understood about the functional and therapeutic implications of this induction^18,19,38^. The multi-gene signature that we uncovered in the atlas using our CoGAPS pattern detection algorithm^36^, associated with inflammatory signaling and EMT in epithelial cells (Pattern 7), was also enriched in our PDO-CAF co-culture, suggesting that the co-culture system can recapitulate fibroblast induction of inflammatory processes in epithelial cells in PDAC. Similarly, EMT has been previously linked to inflammation in PDAC cells and CAFs through an IFNγ response^12^. Applying our established transfer learning methods^33,39^ to our new organoid co-cultures showed, for the first time, that these co-cultures can also model fibroblast-induced changes in epithelial cells associated with inflammation. These data provide supporting evidence that PDO-CAF co-culture can serve as an effective experimental system for future mechanistic studies of these interactions. Specifically, through this analysis we were able to recapitulate the overrepresentation of the inflammatory signal observed in human scRNA-seq data when PDAC PDOs were co-cultured with CAFs, demonstrating that CAFs do alter tumor epithelial cell transcriptional phenotype and overall function. More generally, our combined single-cell analysis and PDO co-cultures suggest that both CAF and epithelial populations are transcriptionally plastic as shown in the changes observed in cell sub-population distribution after co-culture.

Additional studies are needed to distinguish whether fibroblasts directly regulate the inflammatory processes observed in the epithelial cells, whether the inflammatory process is MHC-II-independent or if inflammatory signaling through fibroblasts induces MHC-II expression to a lesser extent than direct activation in response to IFNγ treatment. Still, further ligand-receptor inference analysis of the single-cell datasets uncovered a directed mechanism of signaling between epithelial cells and fibroblasts within human tissue that could be experimentally validated in our organoid co-culture model. Our study identified a fibroblast-induced inflammatory signaling pathway through ITGB1 that was shown to directly influence PDAC cells. While *ITGB1* is a member of the integrin family of proteins that function diversely in cell adhesion and serve as receptors for collagen, it was recently identified as a marker of cytotoxicity potential in CD4+ and CD8+ T cells^40^. Additionally, mutated *ITGB1* was shown to result in an MHC-II-restricted neoantigen in an established sarcoma cell line when evaluated in a subcutaneous xenograft model^41^. Our computational analyses of the single-cell datasets in this study predicted that fibroblasts receive VEGF-A as a ligand to initiate downstream signaling through the VEGF-A pathway that could be experimentally validated in our co-culture PDOs. This finding also supports the inflammatory pattern identified in the tumor epithelial cells in both the atlas and PDO models, as increased *VEGF-A* expression has been identified in alveolar epithelial cells in response to inflammatory stimuli^42^. Further, CAFs were identified in squamous cell carcinoma of the skin as mediators of tumor-enhancing inflammation and angiogenesis, a signature validated in models of mammary and pancreatic tumors^43^. This particular interaction is of biological interest given the role of VEGF-A across different malignancies as well as the ability to inhibit VEGF-A clinically using bevacizumab^44^. While not currently a standard of care agent in PDAC, this finding further demonstrates the clinical relevance of this approach for future mechanistic studies.

While our study robustly demonstrates that computationally inferred intercellular interactions in the TME are preserved between human scRNA-seq datasets and PDO co-culture models, there are also shortcomings to this study. Our collated scRNA-seq atlas of PDAC tumors is restricted to treatment naïve biospecimens from 61 patients, with limited representation of some cell types and limited clinical annotations of the samples. This presents a challenge when trying to relate intercellular dynamics and signaling to patient outcomes for target discovery for translational research. Nonetheless, adapting our validated suite of computational tools to this atlas and the organoid co-culture still provides novel insight into the role of cellular crosstalk in tumorigenesis.

While additional computational tools enable more direct inference of molecular changes from cellular interactions^45^, the unique application of transfer learning between human scRNA-seq data and organoid co-culture enables direct bidirectional investigation of cellular state transitions and intercellular signaling between *in silico* discovery and experimental validation. Currently, this analysis relies on inferences resulting from a pipeline combining NMF-based pattern detection with CoGAPS^36^, transfer learning with ProjectR^33^, and finally ligand-receptor networks from Domino^34^. While CoGAPS and ProjectR allow for unsupervised discovery and query of novel cell states, Domino is limited to investigation of pre-specified pairs of cell types. Additional methods that enable discovery of multicellular interactions and their impacts across both cell types and cellular phenotypes are needed to model the complex processes that underlie carcinogenesis in the PDAC TME. Similarly, while our organoid co-culture model is established to represent the epithelial and fibroblast compartments of the tumor, the immune cells which contribute to the complex TME in this disease are absent in our current PDO co-culture. The data presented here demonstrate the need for immune cell inclusion in future studies, particularly when asking questions related to the tumor immune microenvironment or mechanisms of response or resistance to immunotherapy. With these limitations in mind, we advocate for a complementary approach moving forward that combines reference human single-cell atlases and PDO co-culture to transfer discoveries into mechanistic experiments of the TME effects in PDAC.

In summary, we introduce a novel bidirectional approach leveraging scRNA-seq data and PDO co-culture to examine patterns of inflammation in PDAC. Further, we used this approach to specifically query patterns of inflammation inherent in the malignant epithelial cell compartment, identifying programs of gene expression that are both intrinsic to the epithelial compartment and those that are influenced by tumor residing CAFs. The power of applying computational biology to relate human tissue to organoid co-culture can be exploited in future studies spanning discovery, mechanistic validation, and perturbation of the complex cell-to-cell cross-talk in tumors that underlies tumorigenesis.

## Methods

### scRNA-seq dataset integration and harmonization for the PDAC atlas

The six different datasets provided gene expression data with different versions (GRCh37 or GRCh38) and nomenclatures (Ensembl identifiers vs. HUGO gene nomenclature) of the human reference genome. Available patient metadata are summarized in Table 1. All analyses were performed in R (V 3.6-4.1) or Python (version 3.8). Unified integration of the measured features revealed 15,219 genes that could be matched between all datasets with assured certainty. Next, cells with unfavorable quality, defined as mitochondrial counts >15% and unique features of <50 or >5,000, were removed. Computational pre-processing was performed with the Monocle3^1^ R package. Dimensionality reduction into a unified manifold approximation and projection (UMAP) was based on the first 100 principal components and batch correction was applied per manuscript to account for potential dataset-intrinsic biases (technical or biological) using Batchelor R as utilized by the Monocle3 pipeline^2^. Annotation of cell types is described in detail in the Supplemental Methods. The distributions of epithelial and fibroblast populations and patient-level correlations across epithelial and CAF subtypes are further methodologically detailed and illustrated in Supplemental Fig. 10-12. Plotting was performed with the ggplot2 R package and Excel (Microsoft, Redmond, WA). For high-performance computing tasks, we leveraged the MARCC (Maryland Advanced Research Computing Center, Baltimore, MD) and AWS (Amazon Web Services, Seattle, WA) servers.

### CoGAPs analysis of expression patterns

Non-negative matrix factorization (NMF) of transcript counts was conducted using CoGAPS (V 3.5.8)^3,4^. Given a matrix of single-cell data with normalized expression values, CoGAPS factorizes this matrix into two related matrices of gene weights (amplitude matrix) and sample weights (pattern matrix) for random subsets of the data based on the nsets parameter followed by relearning of the amplitude matrix on the full dataset. CoGAPS was run on log2 transformed counts of 15,176 genes from 25,442 cells in Peng et al and Steele et al annotated as epithelial_normal, epithelial_cancer, or epithelial_unspecified^5,6^.Standard parameters were set to 8 Patterns, 50,000 iterations, seed 367, sparse optimized, and distributed: “Single-Cell”. Sparsity parameters were alpha = 0.01, max Gibb mass 100. Distributed CoGAPS parameters were 15 nSets, cut 10, minNS 8, maxNS 23.

Marker genes for each pattern were identified using the patternMarkers function in CoGAPS (V3.9.5) with the “cut” threshold to provide subsets of the top-ranking genes associated with each pattern^7^. Overrepresentation analysis was then conducted using the fora function in the fgsea R package (V1.18.0) to find enrichment of any hallmark gene sets from the Molecular Signatures Database^8,9^ among the pattern markers for each CoGAPS pattern. The universe used in the overrepresentation analysis was all human genes with HGNC symbols in the GRCh38.p13 genome assembly (n = 39,535)^10,11^.

### Organoid and CAF co-culture and single-cell analysis

Patients with PDAC undergoing endoscopic biopsy or surgical resection were enrolled in IRB-approved tissue acquisition protocols at Johns Hopkins Hospital and Massachusetts General Hospital (MGH) (NCT03563248). Patient-derived organoids (PDOs) were generated from patient surgical specimens following a combination of mechanical and enzymatic dissociation as previously described^11^. CAFs were extracted from surgical resection specimens after straining remnant tissue through a 70μm cell strainer and washed twice with human organoid wash media (Advanced DMEM/F12, 10mM HEPES, 1x GlutaMAX, 100μg/mL Primocin, 0.1% BSA) with centrifugation between washes. For co-culture, organoids were combined with patient-matched CAFs in Matrigel (Corning, 356234) at a 1:10 ratio of organoids to CAFs. In parallel, CAFs and PDOs were plated separately in Matrigel. Co-cultures and monocultures were plated in 24-well tissue culture dishes and extracted after 12 hours using Cell Recovery Solution (Corning, 354253) and incubated on ice at 4°C for 45 minutes for Matrigel depolymerization. Cells were then pelleted and washed in human organoid wash media prior to pelleting again. Organoids were dissociated to single cells using TrypLE Express (ThermoFisher Scientific, 12604013) following manufacturer instructions. Single-cells were barcoded using the MULTI-seq protocol as previously described^12^. Single-cell transcriptomics library prep was completed using the 10x Genomics Chromium Single Cell 3’ Gene Expression Dual Index Library (V3.1) according to manufacturer specifications. Library preparations quality were analyzed using the 2100 Bioanalyzer (Agilent). Sequencing was completed at the Johns Hopkins Genetic Resources Core Facility (GCRF). Cellranger (V6.0.0) was used to generate the feature-barcode matrices, aligned to the hg38 genome. Multiseq10x (V1.0) was used as the preprocessing pipeline companion to split the MULTI-seq FASTQs into cell barcode, unique molecular identifiers (UMI), and sample barcode sequences. Reads that did not align with >1 mismatch to any reference sequence and reads representing duplicated UMIs on a cell-by-cell basis were removed. Demultiplex (V1.0.2) was used for demultiplexing the data. The 3DGE data were log normalized, linear dimension was reduced using principle component analysis, and differentially expressed genes were identified in Seurat by Wilcoxon Rank Sum Test (V4.0.1). Additional annotations of Moffitt classifiers, denoting classical and basal epithelial subtypes, and CAF subtypes were added to the Seurat object metadata based on the clustering^13^. Co-culture cell types were parsed based on these annotations and the barcode distinctions. Projection of the discovered CoGAPS Pattern 7 onto the 12HR MULTI-seq expression data was completed using ProjectR (V1.8.0). The MULTI-seq expression data and CoGAPs feature loadings were run through the projectR function of the package^14^. The projection results were combined with the MULTI-seq metadata and plotted using ggplot2 (V3.3.5) and Wilcoxon results added using ggpubr (V0.4.0)^14^.

### Flow Cytometry and Cell Sorting

Organoids were extracted using Cell Recovery Solution (Corning, 354253) and incubated on ice at 4°C for 45 minutes for Matrigel depolymerization. Cells were pelleted and washed in human organoid wash media. Organoids were dissociated to single cells using TrypLE Express and washed in MACS buffer (PBS + 5 mM EDTA + 1% Fetal bovine serum). Cells were resuspended in PBS + Zombie NIR catalog no. 423106 (dilution 1:1000) + Human TruStain FcX catalog no. 422302 (dilution 1:100) for 10 minutes at room temperature in the dark. Cells were quenched with MACS buffer, spun down, and then resuspended in surface stain for 20 minutes on ice at 4°C in the dark; antibodies were purchased from Biolegend, APC EpCAM catalog no. 324208 (dilution 1:200), PE/Cy7 HLA-A, B, C catalog no. 311429 (dilution 1:200), AF700 HLA-DR catalog no. 307626 (dilution 1:200), PerCP/Cy5.5 PD-L1 catalog no. 329738 (dilution 1:100). Cells were washed twice in MACS buffer. Flow cytometry analyses were performed on the Beckman Coulter Cytoflex.

For co-culture cell sorting, 1mL of 1mg/mL Dispase II (Thermofisher, 17105041) in organoid wash media was added to each coculture and monoculture dome to depolymerize Matrigel for 1 hr at 37°C. Digest was quenched with 1mL of wash media and cells were spun down. Cells resuspended in PBS + Zombie NIR catalog no. 423106 (dilution 1:1000) + Human TruStain FcX catalog no. 422302 (dilution 1:100) for 10 minutes at room temperature in the dark. Cells were quenched with MACS buffer, spun down, and then resuspended in surface stain APC EpCAM catalog no. 324208 (dilution 1:200) and FAP R&D Systems, catalog no FAB3715P-100 (dilution 1:75) on ice at 4°C for 20 minutes in the dark. Cells were washed twice in MACS buffer and filtered through 70um filter. Cell sorting was performed on BD Fusion Sorter.

### Quantitative PCR (qPCR)

To evaluate gene expression of MHC-II genes in patient-derived organoids, total RNA extraction using the RNeasy Mini Kit (Qiagen, - Catalog Number: 74104) was completed for each patient-derived organoid line according to manufacturer specifications. cDNA synthesis was performed using Invitrogen TaqMan Reverse Transcription Reagents (Catalog Number: N8080234), following manufacturer’s instructions. Real-time quantitative PCR was completed using the ThermoFisher Taqman Gene Expression Assays according to manufacturer’s protocol in the QuantStudio 6 Flex System (Applied Biosystems) mRNA targets included: ITGB1 (Hs01127536_m1), VEGFA (Hs00900055_m1), HLA-DRA (Hs00219575_m1), HLA-DRB1 (Hs04192464), HLA-DQB1 (Hs03054971_m1), and HLA-DPB1 (Hs03045105_m1). Relative gene expression was quantified using the 2^−ΔΔCt^ method as previously described^15^, and GAPDH (Hs02786624_g1) was used as the endogenous control. Data were analyzed using Applied Biosystems QuantStudio™ Real Time PCR System Software (V1.7.1).

### Inference of transitions in cellular phenotypes and intercellular interactions

Within each cell group, additional analyses were performed to compute heterogeneity of cellular phenotypes, state transitions, and inter-cellular signaling across the atlas datasets. First, cell cycle scores and phases were computed with tricycle (V1.2.0)^16^. Further unsupervised exploratory analysis of transitions in epithelial cell states was performed with CoGAPS (V3.5.8)^3^ analysis across epithelial populations in tumor and normal samples from Peng et al^5^ and Steele et al^6^. Single cell CoGAPs was run for 8, 10, and 12 patterns. Eight patterns were selected as the final analysis because 12 patterns returned 10 patterns suggesting an overfitting of the data. Further the 8-pattern run resulted in all 8 patterns that were analogous with the other patterns found in the higher dimensional runs. Finally, the impact of fibroblast cells on epithelial cells was computed by estimating intercellular signaling with Domino (V0.1.1)^17^ independently for each of the datasets in the atlas.

For Domino analysis, pyScenic (V0.11.0) for Python was first used to generate the gene regulatory network and co-expression modules, the regulon predictions, and the area under the curve (AUC) matrix of cellular enrichment^18^. This was completed by providing the extracted counts matrix, a list of transcription factors, motif annotations, and cisTarget motifs for the hg38 genome^18^. With the use of the AUC and regulon predictions, a domino object is created and the signaling network built. This allowed for the visualization of global signaling network, gene networks and incoming signaling heatmaps for each narrow subtype annotation, a heatmap of the correlation between transcription factors and receptors, and lastly, the global transcription factor-ligand-receptor network between all subtype annotations^17^.

## Data Availability Statement

Submission of the RNA-seq data to dbGaP is in process. All analysis scripts are available from: https://github.com/fertiglab/

## Acknowledgements

The authors would like to thank Dr. Chris McGinnis for his technical expertise in the completion of the MULTI-seq experiment. We would also like to thank the authors of the 6 datasets used in the generation of the atlas: the laboratories of Dr. Hector Alvarez, Dr. Haiyong Han, Dr. Marina Pasca di Magliano, Dr. David Tuveson, Dr. Wenming Wu, and Dr. Itai Yanai. Sequencing was completed through the Genetic Resources Core Facility, RRID:SCR_018669. Finally, we are grateful to Aviva Fertig for sharing her mommy with us during a pandemic. This work was funded by the Hopper-Belmont Foundation (JWZ), The Lustgarten Foundation (EMJ), Johns Hopkins University Discovery Award (EJF, LW), NIH/NCI (U01CA253403 to EJF; P01CA247886 to EMJ; P30CA006973; R50 CA243627 to LD; K08CA248710 to RAB), Break Through Cancer to LW and EJF, German Research Foundation (KI 2437-2/1 to BKK). TT Seppälä was supported by fellowship grants and research funding from Sigrid Juselius Foundation, Instrumentarium Science Foundation, Emil Aaltonen Foundation, Jane and Aatos Erkko Foundation, Relander Foundation, and the iCAN precision medicine flagship of the Finnish Academy. Stand Up To Cancer–Lustgarten Foundation Pancreatic Cancer Interception Translational Cancer Research Grant (SU2C-AACR-DT26-17 to RAB). Stand Up To Cancer (SU2C) is a division of the Entertainment Industry Foundation and funding is administered by the American Association for Cancer Research, the scientific partner of SU2C). Stand Up To Cancer - Lustgarten Foundation (2015-002 to DTT)

## Author’s Contributions

*B. Kinny-*Köster, *S. Guinn, J*.*A. Tandurella*: data curation, formal analysis, investigation, visualization, writing original draft and review. *J*.*T. Mitchell, D*.*N. Sidiropoulos, M*.*R. Lyman, A*.*B. Pucsek, R. Suri, C. Cherry, L. Danilova, G. Stein-O’Brien*: data curation, investigation, writing and review. *M. Loth, T*.*T. Seppälä, H. Zlomke, J. He, C*.*L. Wolfgang, J. Yu, L. Zheng, D*.*P. Ryan, D*.*T. Ting, A. Kimmelman, A. Gupta, J*.*H. Elisseeff, L*.*D. Wood, L*.*T. Kagohara*: resources, writing review. *E*.*M. Jaffee*: resources, funding acquisition, writing review and editing. *R*.*A. Burkhart:* conceptualization, resources, writing, review and editing. *E*.*J. Fertig, J*.*W. Zimmerman*: conceptualization, data curation, supervision, funding acquisition, formal analysis, visualization, writing original draft, review and editing.

## Supplementary Information

Supplementary Information is available for this paper.

Correspondence and requests for materials should be addressed to Dr. Jacquelyn Zimmerman.

