## Supplemental methods and figures for "Inflammatory Signaling in Pancreatic Cancer Transfers Between a Single-cell RNA Sequencing Atlas and Co-Culture"

**Cell Type Annotations**

Clustering of cells in the unified integrated atlases was generated using the established step-by-step Monocle3 R-package workflow^1^. At resolution 1e-5, clusters were annotated bay using the ‘topmarkers’ function as well as visually evaluating specific gene expression for canonical cell type markers within the generated UMAP **(Supplemental Figure 2)**. The output of the ‘topmarkers’ function gave the top 25 genes differentially expressed by each cluster. Clusters clearly divided the samples by cell type rather than by manuscript or sample. Of note, two of the six datasets (Peng et al^2^ and Elyada et al^3^) provided cell type annotations which we utilized to cross-check the *de novo* cell type annotations.

Epithelial cell clusters were first identified by mapping gene expression of classic epithelial cell markers like *KRT19*, *TSPAN8* and *SLC4A4*. Differential high expression of *SLC4A4* and *AMBP* helped define the benign population of epithelial cells.^2,4,5^ We conclusively defined this population of benign epithelial cells due to this gene expression signature as well as the fact that most cells that cluster here are from control pancreas tissue samples. These cells were confirmed to be benign based on the lack of copy number variations (CNV) found using InferCNV (V1.3.3) **(Supplemental Figure 3)**^6^. Similarly, differential gene expression was used to identify malignant epithelial cell clusters by identifying overexpression of *KRT19*, *TFF2*, *CEACAM6* and *MUC1*^2,4,7^. Cross-checking with provided annotations identified a subset of cells within a malignant epithelial cluster previously annotated as acinar cells by Peng et al^7^. Re-clustering of the respective cluster with higher resolution 1e-4 in Monocle3 resulted in a subset of cells with differentially expressed acinar cell markers (*PRSS1*), which we annotated as acinar cells. We confirmed malignant epithelial cell identity using CNV compared to benign epithelial cells from control pancreas samples. Further classical and basal classification of malignant epithelial cells was performed in PDAC samples with gene sets from Moffitt et al^8^ using the module score function from Seurat^9^. An additional unrelated cluster was annotated as acinar cells according to markers such as *PRSS1* and *CELA3A*^9^.

The fibroblast cluster was initially identified based on differential gene expression of *LUM*, *DCN*, *COL1A1*, *FAP*^10^. When selecting cells of this fibroblast cluster from cancer-derived tissue samples, we refer to these cells as CAFs. Further annotation of inflammatory and myofibroblastic CAFs (iCAFs and myCAFs, respectively) was performed in PDAC samples with gene sets utilized by Elyada et al^3^ using the module score function from Seurat^10^. The stellate cell cluster is defined by *ADIRF*, *RGS5*, and *PDGFRB* expression^7,11^.

We annotated immune cells into 3 separate categories. Immune cells derived from a myeloid lineage were identified by markers such as *S100A9* and *S100A8* (classic neutrophil markers) and *AIF1* (a known marker of macrophages). Lymphoid-derived immune cells were characterized as lymphoid T cells (via *CD3D* and *CD3E* markers) whereas B cells were identified by markers that included *CD79A* and *CD79B*^7,12^. More specifically annotated immune cell populations are included in **Supplemental Figure S2**^2^.

Erythrocytes were identified by markers such as *HBB* and *HBA1*. Endocrine cells (specifically β-cells) were identified using *INS* expression^11^. Finally, the cluster annotated as endothelial cells were identified by the expression of *PLVAP* and *CDH5*^12^.

In total, we annotated 132,136 cells in the complete atlas (94.2% of all cells). The remaining 8,114 cells (5.8%) scattered throughout the UMAP in multiple independent clusters without conclusive differentially expressed marker genes, characterized primarily by patient-specific origin (likely batch-related differences). In the subset of PDAC specimens, we annotated a total of 111,032 cells (98.5% of all cells) while 1,720 cells (1.5%) were not annotated, respectively.

**Determination of mean CoGAPS Pattern 7 weight in epithelial cells in relation to proportion of CAF populations from each patient**

For each sample from the Peng et al^13^ and Steele et al^13^ studies, mean CoGAPS pattern weights among epithelial cells were calculated. Fibroblast proportions were calculated as the number of cells from a sample annotated as “Fibroblast” divided by the total number of cells from that sample in the atlas. iCAF and myCAF proportions were calculated as the number of cells from a sample annotated as “Fibroblasts” and as “iCAF” or “myCAF”, respectively, divided by the number of cells annotated as “Fibroblast.” Linear regression was carried out with the ggplot2 R package (V3.3.5), and Pearson correlation coefficients (R) were calculated with ggpubr (V0.4.0).

**Bulk RNA-seq and analysis**

Total RNA extraction (Qiagen, AllPrep DNA/RNA Mini Kit- Catalog Number 80204) was completed for 13 patient-derived organoid lines according to manufacturer specifications. Total RNA was then quantified using the Qubit RNA High Sensitivity Assay kit (ThermoFisher Scientific). Bulk RNA-seq was performed through the Johns Hopkins Experimental and Computational Genomics Core (ECGC). 500 nanograms of total RNA was used in the Illumina TruSeq Stranded Total RNA LT (Illumina, San Diego CA; Cat. No. RS-122-2201, RS-122-2202, and RS-122-2203) approach, which utilizes ribosomal RNA depletion and polyadenylated RNA enrichment selection, according to the manufacturer’s recommendations.  Quality and quantity of the resulting cDNA libraries was monitored using the Bioanalyzer High Sensitivity kit (Agilent Cat. No. / ID: 5067-4626).  mRNA libraries were sequenced on an Illumina Novaseq 6000 (Illumina, San Diego CA) instrument using 150bp paired-end dual indexed reads and 1% of PhiX control to a target depth of approximately 50,000,000 reads per sample. Reads were aligned to the hg38 reference genome using rsem (V1.3.0) with the following options: star-calc-ci-star-output-genome-bam-forward-prob 0.5. Differential expression analysis and statistical testing was performed using DESeq2 R/Bioconductor package^14^.

CibersortX was used to deconvolute the basal and classical subtypes annotated with the atlas: Peng Epithelial cells onto the Bulk data^15^. A signature matrix was generated from the scRNA-seq data reference file (Atlas Peng Epithelial). The mixture file was created using the Bulk RNA-seq data. Both sets of data are in non-log space prior to the creation of the signature matrix and mixture files. Cross-platform deconvolution was completed using the single cell signature matrix file and Bulk RNA mixture file with s-mode batch correction and 100 permutations. These results were then added to the DESeq object used to store the bulk data as an additional sample annotation. All plots generated were completed using ggplot2 (V3.3.5) in R (V4.1.1). DESeq object was designed using the classical subtype annotations gathered from the CibersortX results.

**Immunohistochemistry (IHC)**

Patient-derived organoid and co-culture samples were fixed in 10% formalin and embedded in paraffin at the Sidney Kimmel Comprehensive Cancer Center Oncology Tissue Services Core. Automated staining was performed on 4µm sections using the Leica Bond RX system (Leica Biosystems, Buffalo Grove, IL) and the Bond Polymer Refine Kit (DS9800, Leica Biosystems, Buffalo Grove, IL). Slides were baked online followed by low pH antigen retrieval. Endogenous peroxidase was blocked using Peroxide Block (DS9800, Leica Biosystems). Anti-HLA-DR clone EPR3692 (ab92511, abcam plc., Cambridge, MA) or anti- HLA-DR/DP/DQ clone CR3/43 (M077501-2, Agilent Technologies Inc., Santa Clara, CA) antibodies were applied for 15 min at room temperature using a concentration of 0.789 µg/mL or 0.077 µg/mL, respectively, using Antibody Diluent (S302283-2, Agilent Technologies Inc). Detection was performed using the Bond Polymer Refine Kit (DS9800, Leica Biosystems). Slides were counterstained and coverslipped using Ecomount (5082832, Biocare Medical, Walnut Creek, CA). Slides were imaged using a Hamamatsu NanoZoomer digital slide scanner at 20x magnification using the associated NDP.toolkit slide processing software.

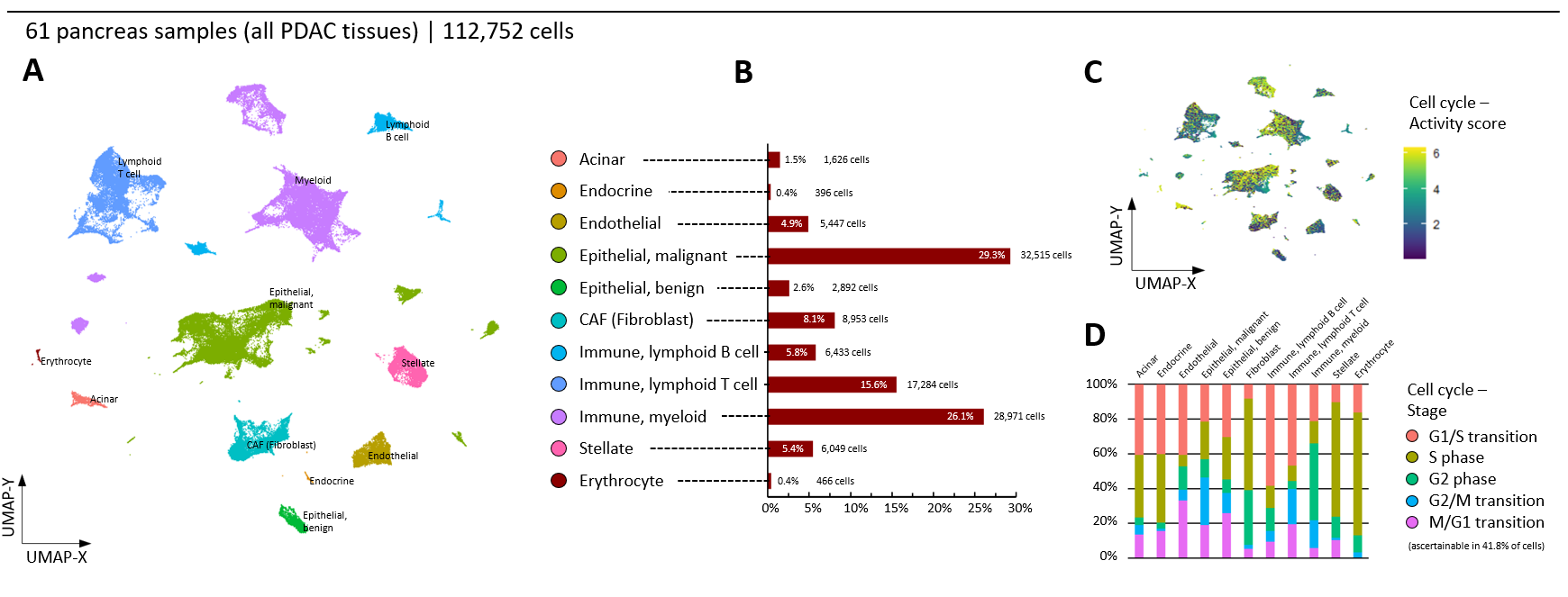
**Supplemental Figure 1.** (A) Complete atlas subset of tumor specimens with assigned cell types and (B) overall cell type composition. (C) Cell mapping of cell cycle activity scores and (D) cell cycle phases in atlas samples derived from tumor tissue.

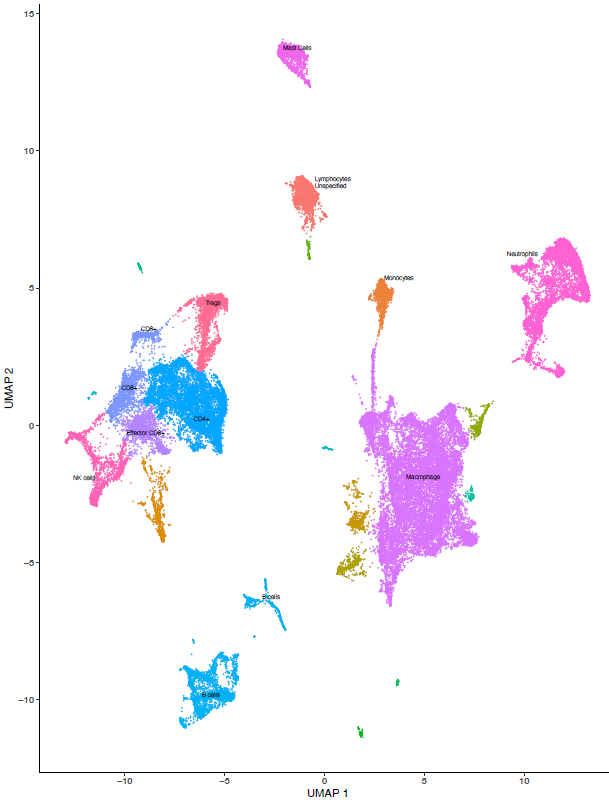

**Supplemental Figure 2.** Subpopulation annotation of immune cells included in the atlas.

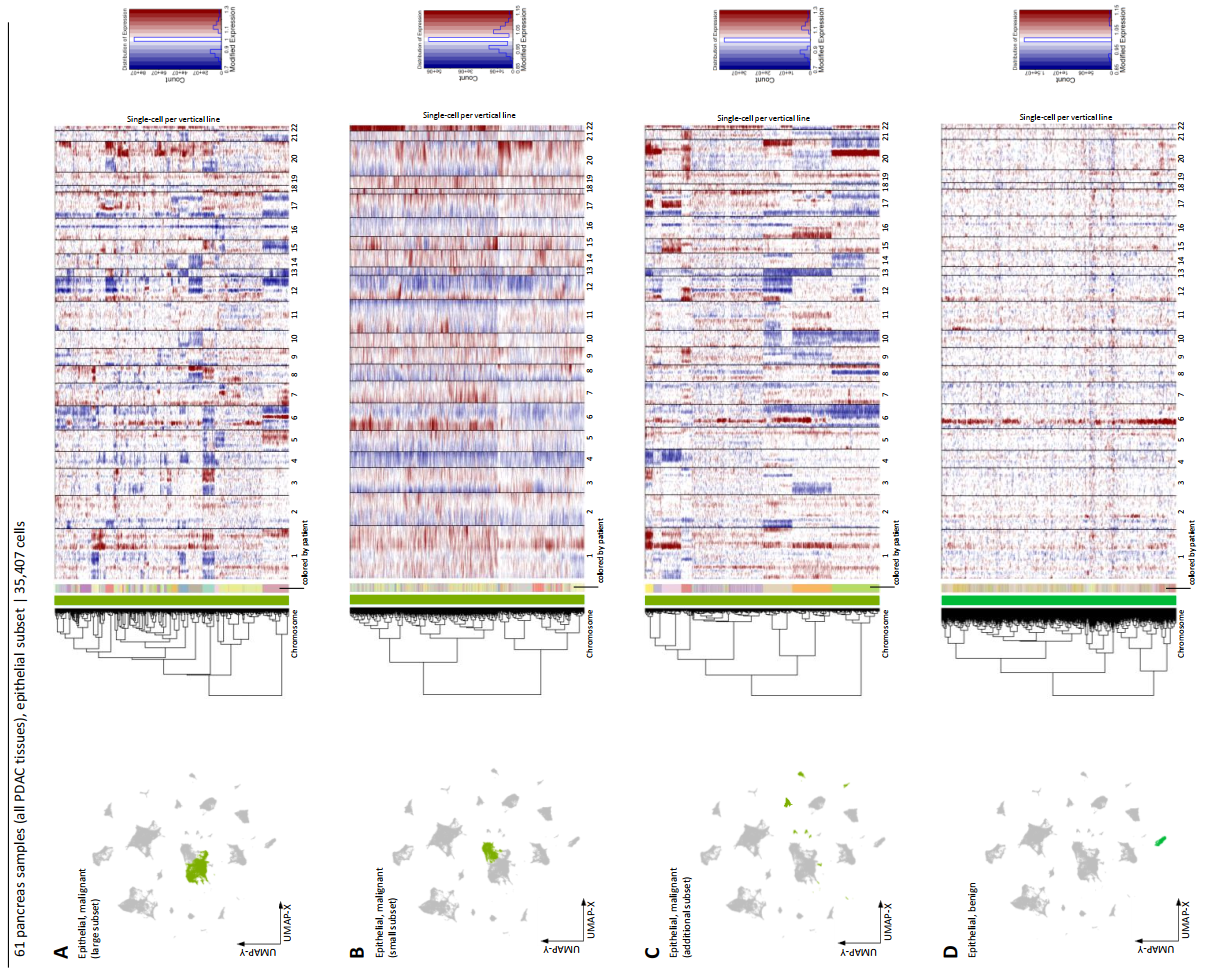

**Supplemental Figure 3.** Copy number variation analyses for epithelial cell populations from PDAC pancreas tissue samples derived by InferCNV. A random subset of 1,000 epithelial cells derived from 16 non-malignant pancreas tissue samples served as a reference for copy numbers. The analyses confirm that the cell genomes selected in (A), (B), and (C) are malignant and that cells in (D) resemble the reference subset indicating benign genomes (homogenous amplification in chromosome 6 across all included patients, interpreted to be likely a batch effect).

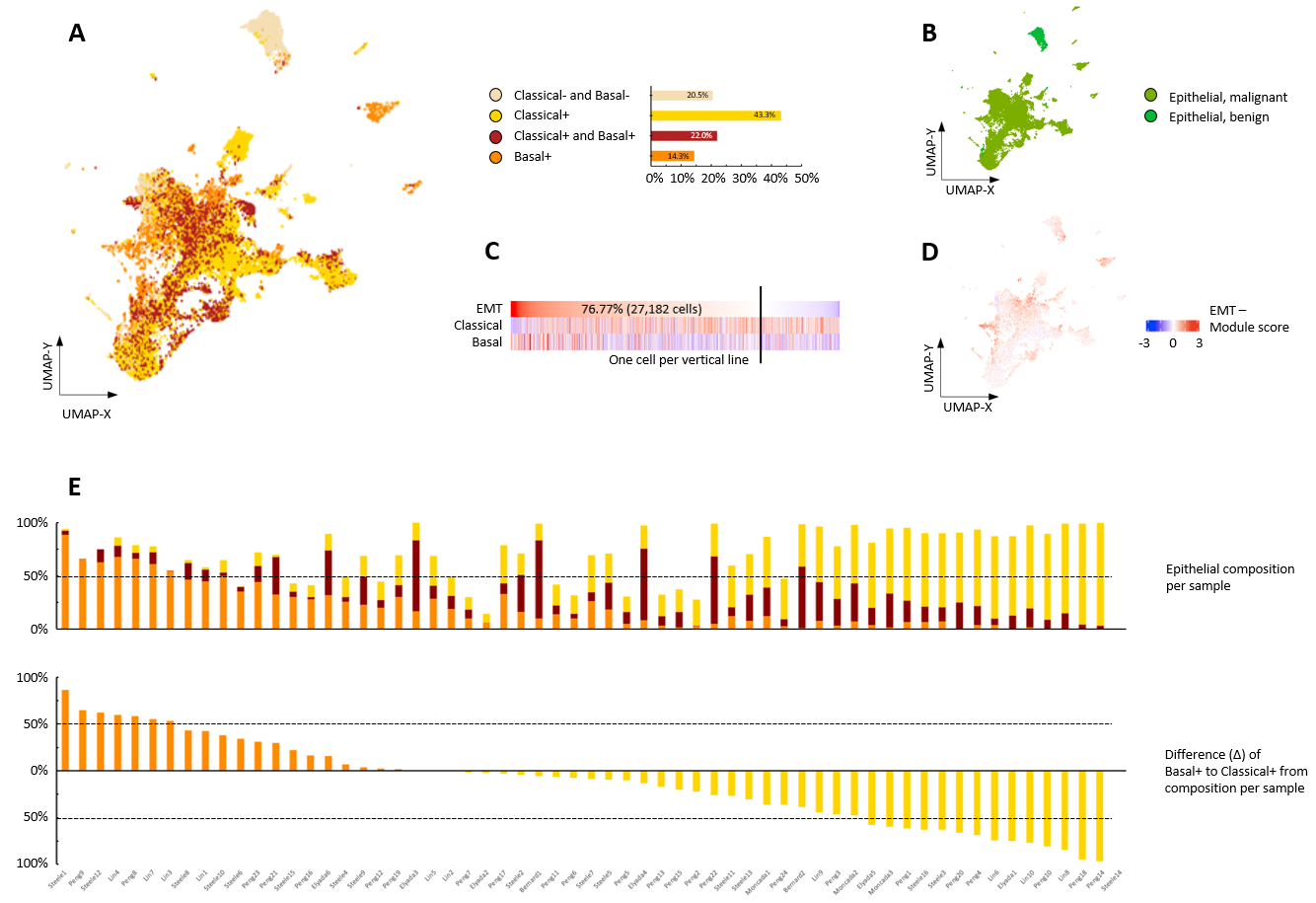

**Supplemental Figure 4.** Selection of epithelial populations from the atlas after subsetting. (A) Cell mapping based on gene set expression scores for Classical and Basal subtypes according to the Moffitt classification and subtype composition according to four classifiers. We leveraged the ‘module score’ function from the Seurat R package to calculate comparable (scaled) scores per cell for the summarized expression of supervised Classical and Basal gene sets^8,9^. Overall, 65.2% of the cells had a positive Classical and 36.3% a positive Basal score. However, not all cells distinctly aligned with one of these subgroups. Within the total epithelial cell population, 22.0% of the cells had overlapping (dual) positivity and 20.5% dual negativity for the two classifiers. Of the malignant epithelial cells, 70.7% demonstrated a positive Classical score and 40.0% a positive Basal score. Moreover, 23.9% of the malignant cells showed dual positivity and 13.2% of cells were negative for both Classical and Basal subtypes, while 46.8% demonstrated a discriminative Classical and 16.1% a discriminative Basal expression pattern. Visually, in the two-dimensional atlas, the population of dual positive cells localized between cells with distinctly Classical and Basal properties. (B) Cell mapping of the initial cell type annotations from the PDAC-excl atlas in the epithelial cells from tumor pancreas tissue samples. (C) Heatmap of epithelial-to-mesenchymal transition (EMT), Classical and Basal gene set scores sorted by EMT score. 76.8% of the epithelial population displayed a positive EMT score^15^. A high EMT score was correlated with high Basal expression scores (*r* = 0.33, p<0.001), whereas the Classical expression program was inversely correlated with EMT (*r* = -0.28, p<0.001). (D) Cell mapping based on quantitative EMT gene set score in epithelial cells from tumor pancreas tissue samples. (E) Patient-level composition of epithelial classifiers and difference between Basal to Classical fraction, both sorted by difference as depicted in legend below. Although every patient had epithelial cells with a positive Classical and Basal subtype score, most patients had a predominance for either of these scores. Hence, 20 of the 60 patients with scalable scores had a predominant Basal population whereas 37 patients had a predominant Classical population. However, in 10 patients the dual-positive fraction of cells was larger than the distinct Classical- or Basal-positive cells. Of note, in 25% of the samples the dual-negative fraction consisted of ≥50% of epithelial cells demonstrating a dominating functional program not captured by either Classical or Basal properties. One sample did not contribute >3 cells to the assigned epithelial populations and was excluded from patient-level plotting.

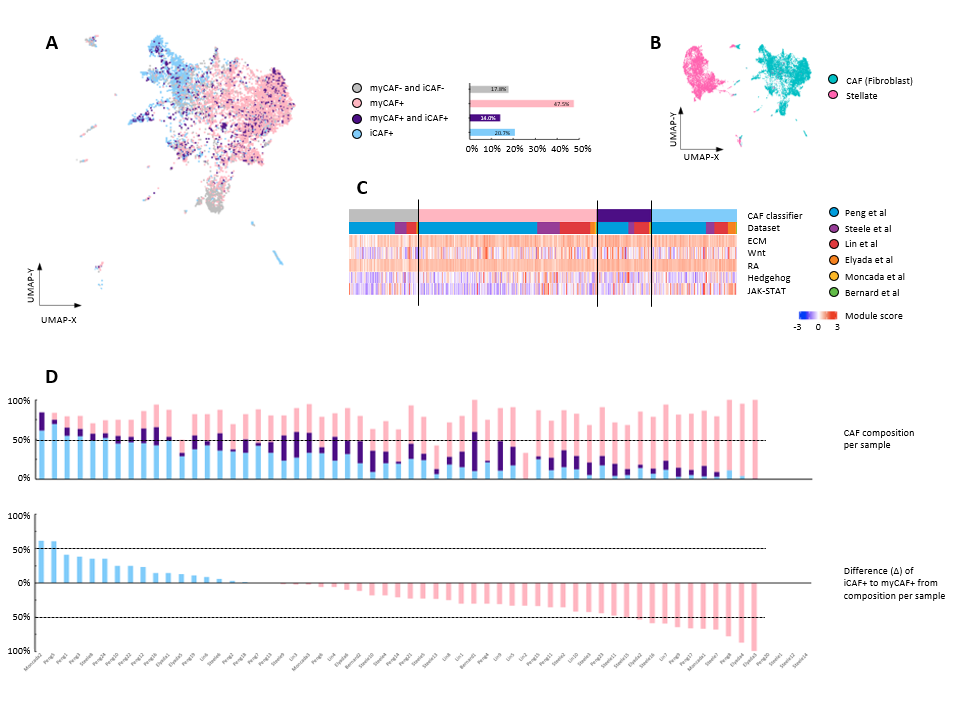

**Supplemental Figure 5.** Selection of mesenchymal populations, CAF and stellate cells, from the PDAC tumor samples atlas. 15,002 cells were used for this analysis. (A) Cell mapping of CAFs based on gene set expression scores for myofibroblastic (my) and inflammatory (i)CAFs and subtype composition according to four classifiers. (B) Cell mapping of the initial cell type annotations from the PDAC-excl atlas in the mesenchymal cells from tumor pancreas tissue samples. For the CAF population, we calculated gene set scores with published features^3^ of the myofibroblastic (myCAF; *ACTA2*, *TAGLN*, *MYL9*, *TPM2*, *MMP11*, *POSTN*, *HOPX*, *TWIST1*, *SOX4*) and inflammatory (iCAF; *CXCL1*, *CXCL2*, *CCL2*, *CXCL12*, *PDGFRA*, *CFD*, *LMNA*, *DPT*, *HAS1*, *HAS2*) expression programs. Of the 8,953 CAFs, 60.1% demonstrated a positive myCAF score and 36.0% a positive iCAF score. Dual positive or negative overlap occurred in 14.0% and 17.8% of the CAFs, respectively. In contrast, 46.1% of the cells demonstrated a discriminative myCAF and 22.1% an iCAF expression program. When analyzing the UMAP visually, cells with either distinct myCAF or iCAF properties clustered closely together with a clear border between the two groups (dual positive cells scattered throughout). This indicates that features of the supervised myCAF and iCAF gene sets were part of the principal components that decomposed variance and informed dimensionality reduction. Scoring for myCAF and iCAF properties at the cell-level also allowed for a comparative analysis of pathway activities based on gene expression scores. (C) Heatmap of extracellular matrix remodeling (ECM), Wnt pathway, retinoid acid pathway (RA), Sonic-Hedgehog pathway (Hedgehog) and JAK-STAT pathway gene set scores sorted by CAF classifier (primarily) and dataset origin. No differences were observed between iCAFs and myCAFs when examining extracellular matrix remodeling (ECM), Wnt and retinoic acid pathways; however, myCAFs showed increased activity for the Sonic-Hedgehog pathway and decreased activity of the JAK-STAT pathway when compared with iCAFs (D) Patient-level composition of CAF classifiers and difference between iCAF to myCAF fraction. 17 of the 57 patients with calculable scores had a predominant iCAF population, and 38 patients possessed a predominant myCAF population. In patients with either a high predominance for iCAFs or myCAFs, the dual-positive fraction of cells was small indicating a polarization towards one of the two gene expression programs within patients. Furthermore, the dual-negative fraction consisted of ≥50% of cells only in three patients suggesting that the majority of functional properties were captured by myCAF or iCAF function Four samples did not contribute >3 cells to the assigned CAF population and were excluded from patient-level plotting. Besides our myCAF/iCAF focus, during initial clustering of the stromal cells from PDAC tumors we also observed an additional population of 273 cells in the atlas (79.9% annotated as stellate cells, 20.1% as CAFs) that expressed HLA genes. Biologically, this cluster reproduced the antigen-presenting CAF (apCAF) phenotype described by Elyada et al^3^. Altogether, this cluster captures the expression program in 1.8% of the mesenchymal cells with contributions from 4 different datasets and 54.2% of PDAC tumors confirming the broader presence of apCAFs across sample cohorts.

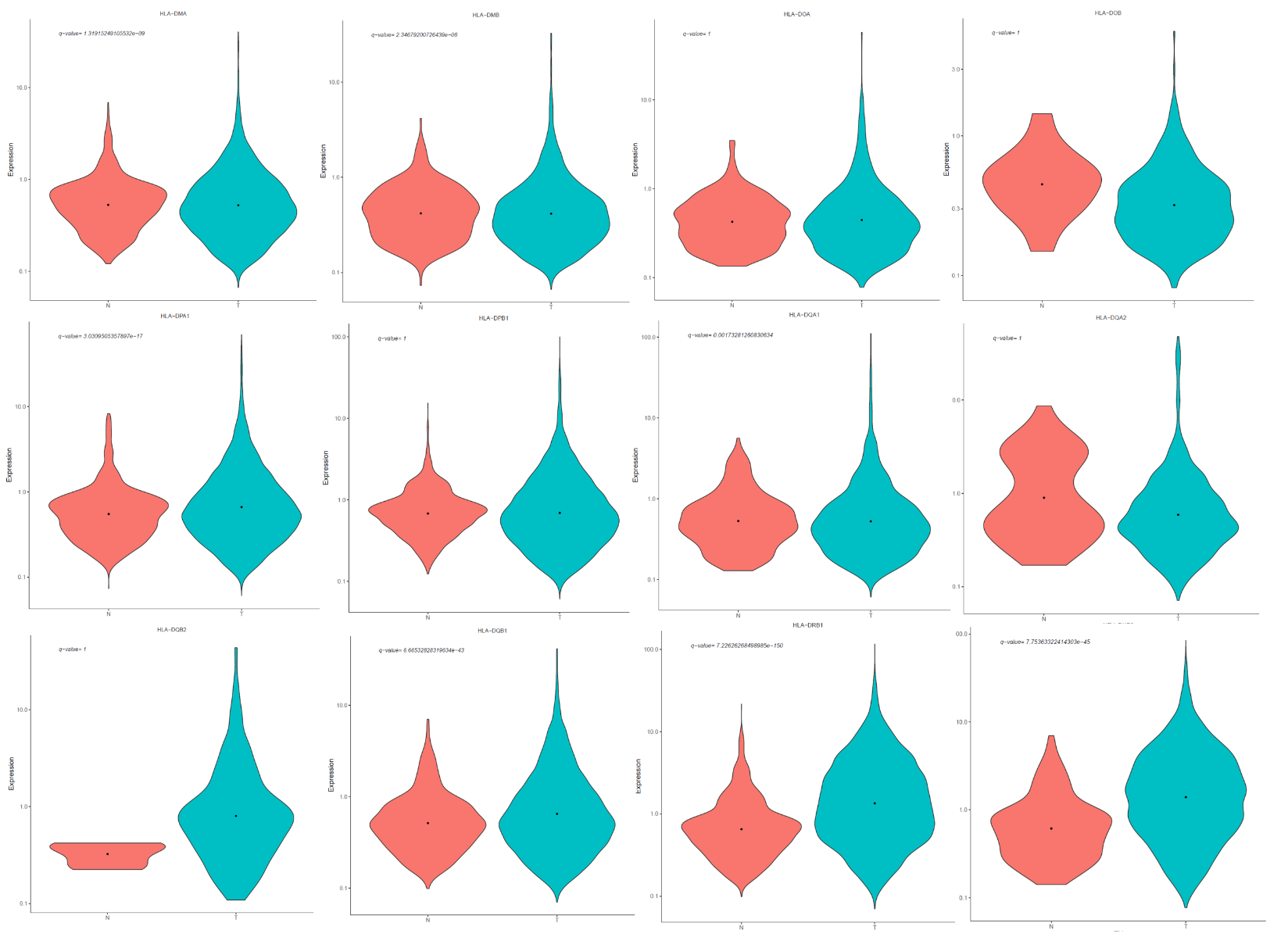

**Supplemental Figure 6.** Violin plots demonstrating differential expression of 12 MHC-II genes in the epithelial cells from atlas samples. Plotted are the single cell raw counts for the 12 MHCII related HLA genes for our Tumor (T, teal-colored) and Normal Control (C, salmon-colored) Samples using Monocle3^1^. Q-value above has been calculated using FDR from differential gene expression of the whole genome using quasipoisson. As shown, expression of HLA-DMA, HLA-DMB, HLA-DPA1, HLA-DQA1, HLA-DQB1, HLA-DRB1, HLA-DRB5 are up-regulated in the Tumor samples with Q-values < 0.01 compared to that of the normal controls.

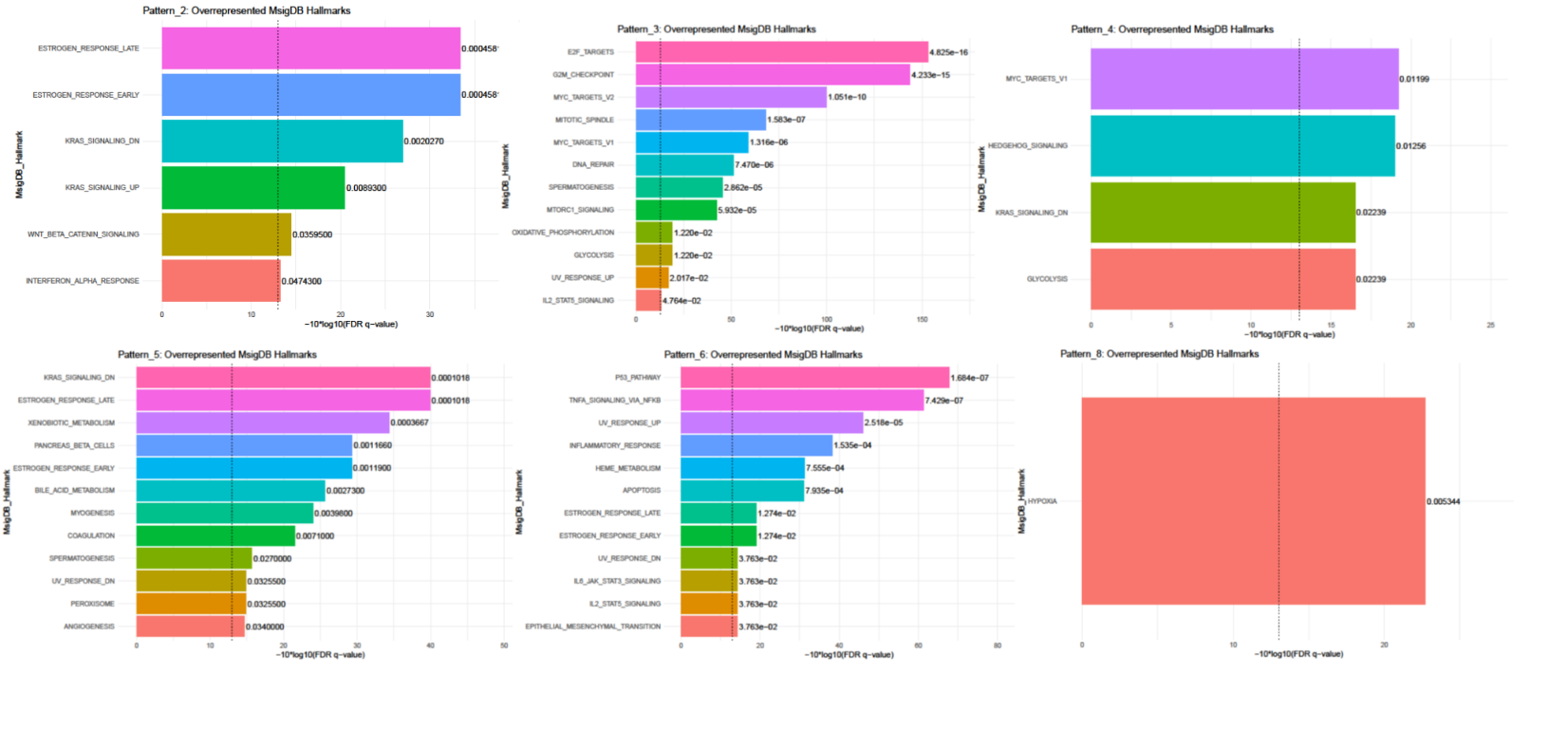
**Supplemental Figure 7.** CoGAPS results evaluating patterns of gene expression across epithelial populations. Pattern 1 did not reach statistical significance; gene sets are included in Supplemental Table 2.

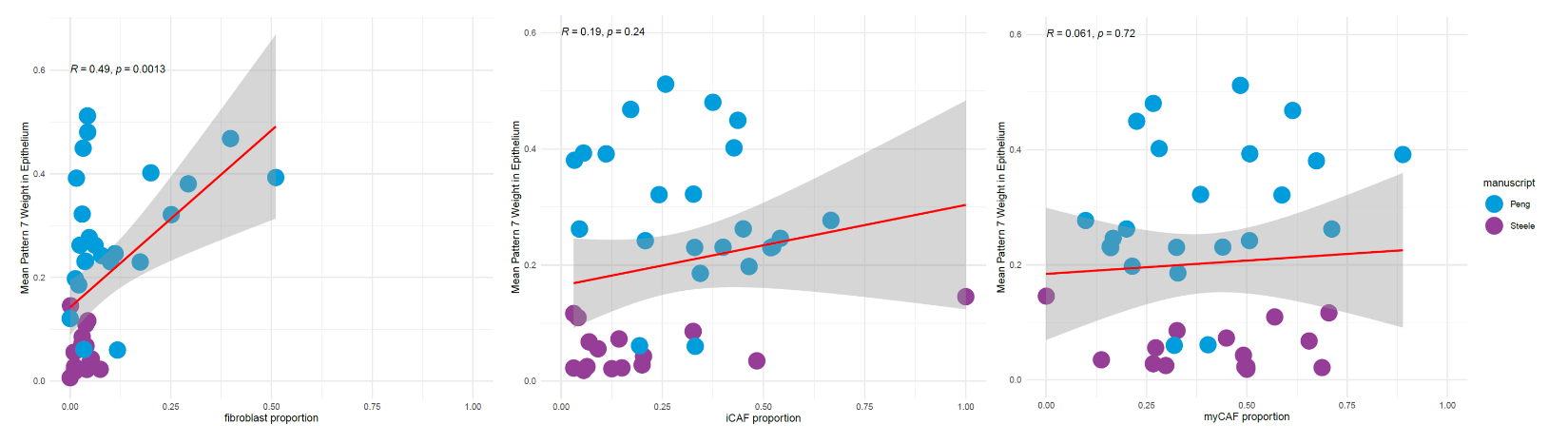

**Supplemental Figure 8.** Correlation of mean pattern weight of Pattern 7 in epithelial cells as it relates to proportion of CAFs per patient. (Left) There is a significant association between total fibroblast content per sample and mean weight of Pattern 7. When fibroblast population is subdivided by iCAF (middle) and myCAF (right), this significant correlation is lost. Each blue circle represents a patient included in the Peng et al^2^ study, and each purple circle represents a patient included in the Steele et al^13^ study.

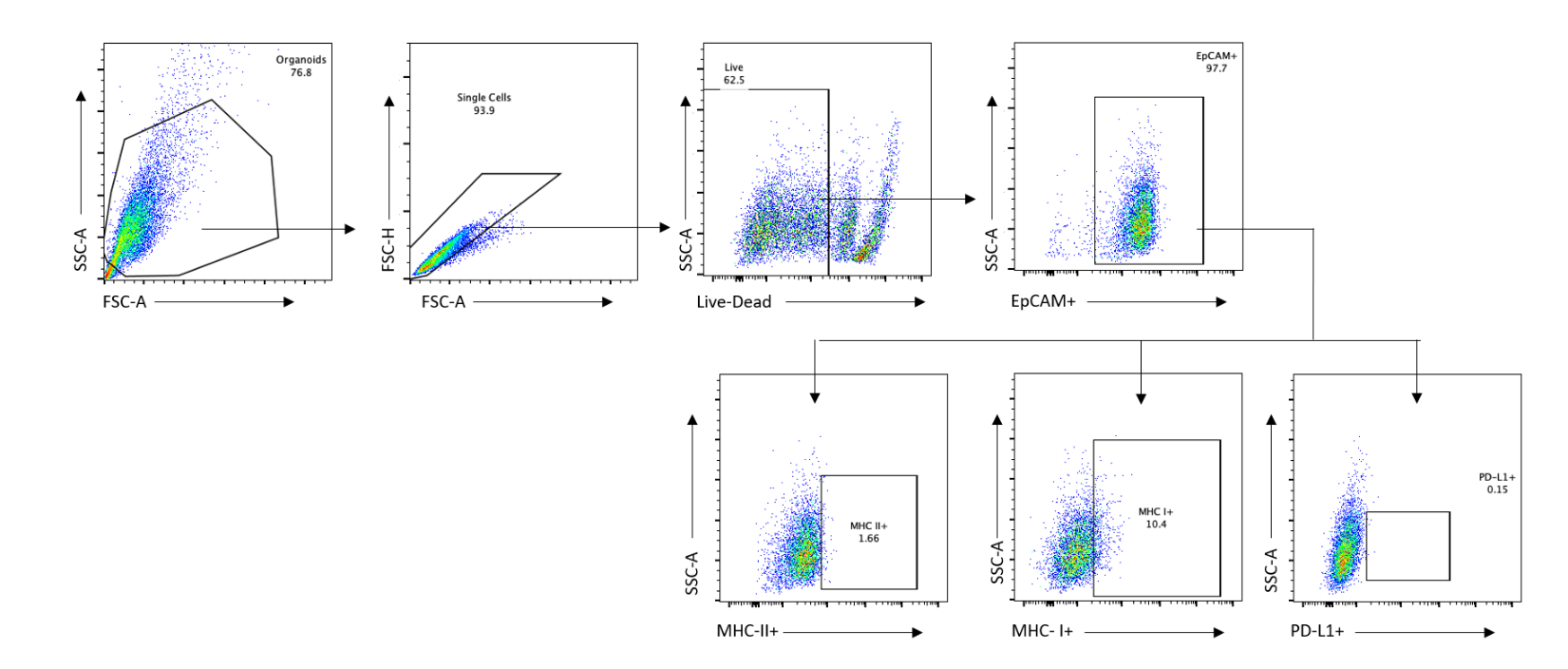

**Supplemental Figure 9.** Representative flow cytometry gating strategy for IFN𝛾 treated organoids. JHH616 p15 untreated at 24 hours shown above.

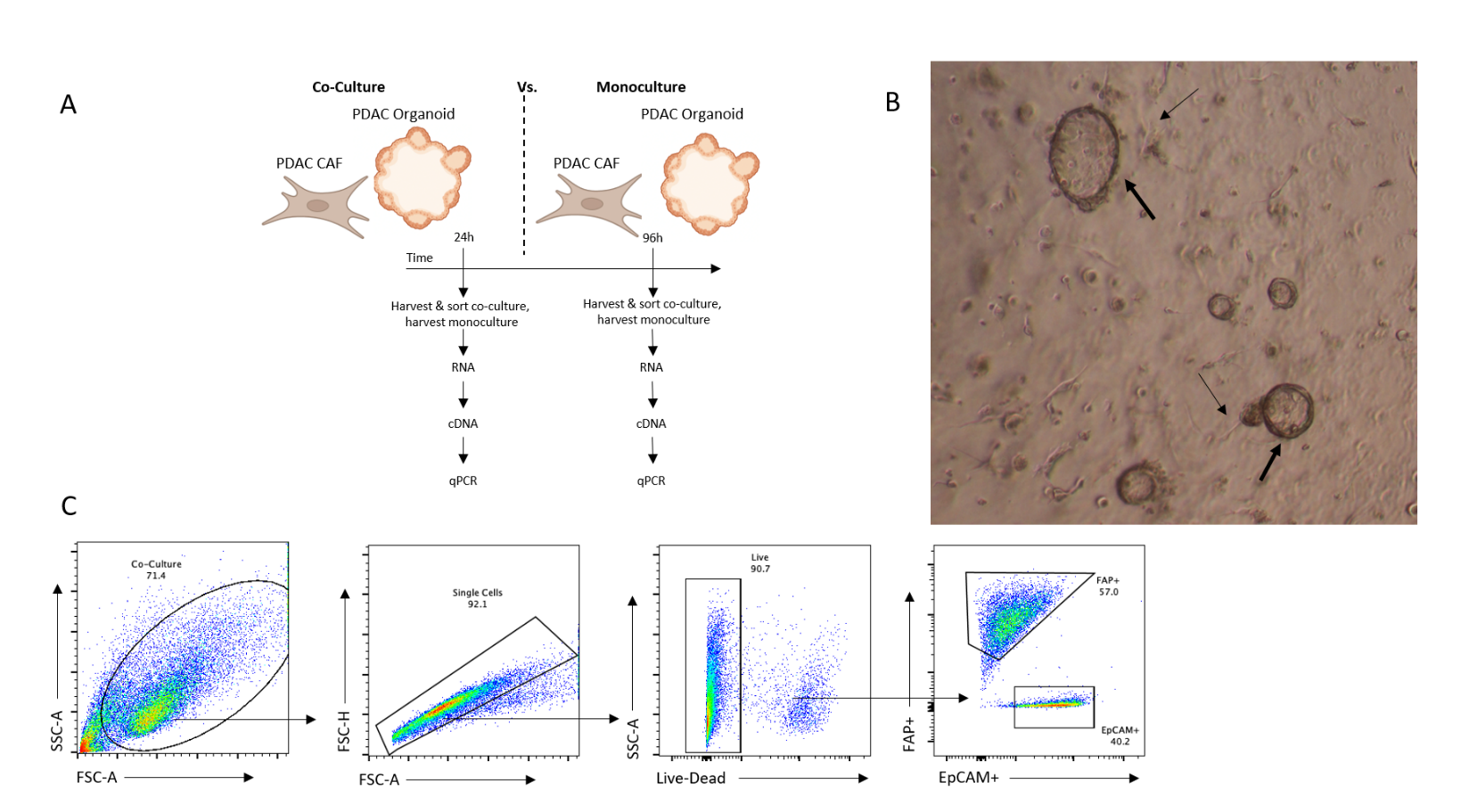

**Supplemental Figure 10.** Optimization of co-culture conditions and viability at time of extraction from culture. (A) Experimental Schematic of coculture set up. (B)Representative co-culture image showing intermingling of organoids (thick arrow) and fibroblasts (thin arrow). (C) Flow plots from co-culture sort. Cells were then processed for RNA in preparation for PCR.

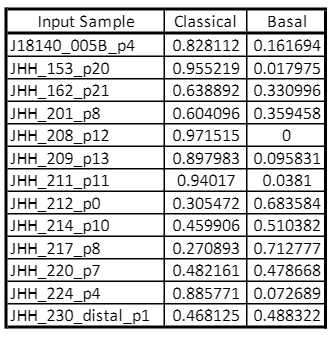

**Supplemental Figure 11.** Heterogeneity of epithelial cells in PDOs. Table of the contribution of the basal and classical programs of epithelial cells in 13 PDO lines following bulk sequencing and subsequent projection onto the atlas.

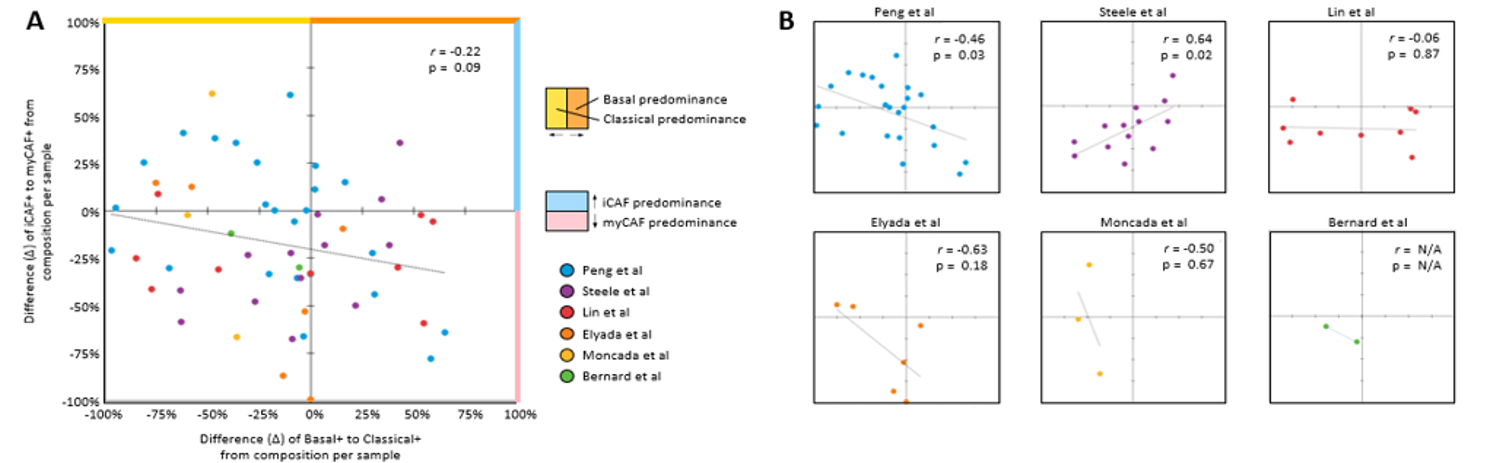
**Supplemental Figure 12.** Patient-level correlation between epithelial subtype composition (difference between Basal to Classical fraction) and CAF subtype composition (difference between iCAF to myCAF fraction) within tumor tissue samples combined (A) and per dataset (B). Correlation coefficients represent Pearson statistic. Despite trends in individual datasets, there was not an association across the patient samples as a whole. This may be due to lack of robust association or sampling error given the relatively low number of CAFs represented in the datasets relative to their contribution to the TME.
